# A dose-sensitive OGT-TET3 complex is necessary for normal Xist RNA distribution and function

**DOI:** 10.1101/2021.06.19.449113

**Authors:** Elizabeth Allene Martin, Jason C. Maynard, Joel Hrit, Katherine Augspurger, Colette L. Picard, Suhua Feng, Steven E. Jacobsen, Alma L. Burlingame, Barbara Panning

## Abstract

Female (XX) mouse embryonic stem cells (mESCs) differ from their male (XY) counterparts because they have lower levels of 5-methylcytosine (5mC) and 5-hydroxymethylcytosine (5hmC). This difference in DNA modifications is a result of having two X chromosomes (Xs), both of which are active at this developmental stage. We identified an X-linked gene, *Ogt*, that controls levels of 5mC and 5hmC in mESCs. OGT is a post-translational modification enzyme and we identified the 5-methylcytosine dioxygenase TET3 as an OGT target that is differentially modified in XX and XY mESCs. In addition to influencing 5mC and 5hmC abundance, OGT dose also controls TET3 and OGT distribution. OGT and TET3 are predominantly nuclear in XX mESCs and cytoplasmic in XY mESCs. Furthermore, these proteins are present in different complexes in XX and XY mESCs. Mutational analysis revealed that TET3 determines the XX-specific abundance of 5mC and 5hmC in mESCs. While TET3 null XX mESCs exhibited modest changes in gene expression, there were substantial alterations upon differentiation into epiblast-like cells (mEpiLCs). In addition, these TET3 null XX mESCs did not undergo X-chromosome inactivation (XCI) when differentiated. These data suggest that an X-dose sensitive complex containing OGT and TET3 regulates cytosine modifications and XCI.

## Introduction

Female (XX) mouse embryonic stem cells (mESCs) exhibit lower global 5-methyl cytosine (5mC) and 5-hydroxymethylcytosine (5hmC) than male (XY) mESCs (Habibi et al., 2013, Zvetkova et al., 2005). These differences have been attributed to X-chromosome (X) dosage, since XX mESCs that have lost one X (XO), display XY-like cytosine modification levels (Zvetkova et al., 2005, Schulz et al., 2014). Both Xs are active in XX mESCs and these cells therefore have a higher dose of X-linked transcripts than XY or XO mESCs (Song et al., 2019), suggesting that one or more X-linked gene products regulate 5mC and 5hmC levels. One X-linked gene implicated in regulation of cytosine modifications is *O*-linked N-acetylglucosamine (*O*-GlcNAc) Transferase (OGT) (Lewis et al., 2014).

OGT is the sole enzyme responsible for attaching a single *O*-linked GlcNAc sugar to serine and threonine residues on thousands of nuclear, cytoplasmic, and mitochondrial proteins (Haltiwanger et al., 1990; Hanover et al., 2012; Maynard et al., 2016). Like phosphorylation, *O*-GlcNAcylation is a reversible modification that affects the function of target proteins. OGT has been linked to regulation of cytosine modifications because it stably interacts with and modifies the Ten-Eleven Translocation (TET) family of enzymes (Vella et al., 2013; Deplus et al., 2013; Chen et al., 2013). The three TETs (TET1, TET2, TET3) are dioxygenases that each oxidize 5mC to 5hmC. 5hmC is not recognized by the maintenance DNA methyltransferase (DNMT1), resulting in passive demethylation during replication (Parry et al., 2021). TETs also oxidize 5hmC to 5-formylcytosine and 5-carboxylcytosine, which are substrates for base excision repair (Tahiliani et al., 2009; Ito et al., 2010; Kriaucionis and Heintz, 2009; He et al., 2011; Ito et al., 2011). Because these higher oxidized states are replaced by cytosine during repair, TETs can also direct active demethylation that does not rely on DNA replication (Ross et al., 2019; Parry et al., 2021).

OGT’s interaction with and modification of TETs can stimulate TET activity (Hrit et al., 2018) and can control TET nucleocytoplasmic distribution (Zhang et al., 2014), providing two mechanisms by which this post-translational modification enzyme could regulate TETs and thus cytosine modifications. To test whether OGT is one of the X-linked proteins that regulates 5mC and 5hmC in mESCs, we manipulated OGT dose in XX and XY mESCs and found that OGT abundance controls cytosine modifications. Our search for OGT targets that are differentially modified revealed that TET3 is more modified in XX than in XY mESCs. In addition, TET3 differs in abundance and distribution and forms distinct complexes in these two cell types. OGT dose also controls TET3 and OGT nucleocytoplasmic distribution. We found that in TET3 null XX mESCs 5mC and 5hmC abundance is altered without notably affecting gene expression. To investigate the developmental significance of TET3, we examined the effects of this null mutation on the ability of cells to undergo X chromosome inactivation (XCI), an epigenetic change that occurs when XX mESCs are differentiated into the next developmental stage, epiblast-like (mEpiL) cells. The establishment of XCI is characterized by the up-regulation of a non-coding RNA, Xist RNA, which remains in the nucleus and ‘coats’ the X concomitant with silencing (Boeren et al., 2021). In TET3 mutant XX mEpiLCs, Xist RNA exhibits abnormal distribution and silencing defects. These results link the activity of a dose-sensitive complex containing X and autosomal proteins to regulation of cytosine modifications and XCI.

## Materials and Methods

### Mouse embryonic stem cells (mESCs) and culture conditions

mESCs (Supplemental methods 1A) were routinely passaged by standard methods using serum+LIF (mESC) media consisting of KO-DMEM, 10% FBS, 2 mM L-glutamine, 1X non-essential amino acids, 0.1 mM 2-mercaptoethanol and recombinant leukemia inhibitory factor. Cells were also grown in 2i media consisting of N2B27 media (DMEM/F12, neurobasal media, 2mM L-glutamine, 0.1mM 2-mercaptoethanol, 100X N2 supplement, 50X B27 supplement) and supplemented with 3mM CHIR99021, 1mM PD0325901, and recombinant leukemia inhibitory factor. To achieve EpiLC differentiation, tissue-culture dishes were incubated with Geltrex (Gibco) diluted in DMEM/F12 media for 30 minutes and replaced with N2B27 media supplemented with 10ng/mL FGF2 and 20ng/mL Activin A (mEpiLC media). The cells were cultured in mEpiLC media for five days, replacing the media every day.

### CRISPR/Cas9 Gene Editing

XX WT/OGT-deg and XX-OGT-GFP mESCs were derived from LF2 XX mESCs using homologous-directed repair CRISPR/Cas9 genome editing. A guide RNA to the OGT 3’UTR (Supplemental methods 1B) was cloned into the px459-Cas9-2A puromycin plasmid using published protocols (Ran et al., 2013). Templates for homology directed repair were amplified from gene blocks (IDT) (Supplemental methods 2-3). Plasmid and template were co-transfected into LF2 mESCs using FuGENE HD (Promega) according to the manufacturer’s protocol. After two days, cells were selected with puromycin for 48 hours, then allowed to grow in antibiotic-free media. Cells were monitored for green fluorescence and single fluorescent cells were isolated by FACS. All cell lines were propagated from a single cell and correct insertion was confirmed by PCR genotyping (Supplemental methods 3).

XX-TET3KO mESCs were derived from LF2 XX mESCs using non-homologous end joining CRISPR/Cas9 genome editing. Two guide RNAs were selected approximately 700 base pairs apart in the first coding exon, exon 3, of *Tet3* (Supplemental methods 1B). They were individually cloned into the px458-Cas9-2A-GFP plasmid using published protocols (Ran et al., 2013). Plasmids were co-transfected into LF2 mESCs using FuGENE HD (Promega) according to the manufacturer’s protocol. After two days, single GFP-positive cells were isolated by FACs and placed into individual wells of 96-well tissue-culture plates. All cell lines were propagated from a single cell and TET3 knockouts identified by immunoblotting. *Tet3* disruption was confirmed by sequencing of each allele (Supplemental methods 3; Fig 4.-fig. sup. 1-3), which identified indels and point mutations. Chromosome number of all cell types was determined by counting metaphase spreads (LF2, E14, LF2-XO) or by G-banding (LF2, XX WT/OGT-deg lines, XX-TET3KO lines).

### DNA cytosine modification immunofluorescence staining

For mESCs, cell pellets were resuspended and incubated for 10min in 0.075 M KCl hypotonic solution. After removing the hypotonic solution by centrifugation, a fixative solution (3:1, methanol:glacial acetic acid) was added dropwise to the cell suspension. Cells were collected, rinsed with fixative solution at least twice, and dropped onto glass slides. For mEpiLCs, cells were cytospun onto glass slides at 800rpm for 3 minutes and then fixed in methanol:acetic acid (3:1) solution for 5 minutes and washed with PBST (0.01%Tween in PBS) for 5 minutes.

To denature DNA, the fixed cells were washed twice with PBST and incubated in 1N HCl at 37°C for 30 minutes. The HCl was neutralized in 100mM Tris pH 7.6 at room temperature for 10 minutes, and then the cells were washed twice more with PBST. The slides were blocked in 5% goat serum, 0.2% fish skin gelatin, and 0.2%Tween in PBS (IF blocking buffer) at room temperature for one hour. Primary antibodies against 5mC and 5hmC (Supplemental methods 1C) were incubated on cells for one hour at room temperature. Then cells were washed twice with PBST and incubated with secondary antibodies (Jackson Immunoresearch) for one hour at room temperature. After incubation, slides were washed with PBST three times. On the second wash, DAPI was added to the PBST. The cells were then mounted using prolong gold antifade (Molecular Probes).

### DNA cytosine modification fluorescent intensity image quantification

Nuclei were masked and pixel intensity determined using ImageJ. The pixel mean intensity value was used to quantify fluorescent intensity inside each nucleus. 50 cells were quantified per condition. Significance was determined by using the “paired two samples for means” t-test.

### Protein immunofluorescence staining

mESCs were cytospun onto glass slides at 800rpm for 3 minutes and then fixed in 4% paraformaldehyde for 10 minutes and washed with PBST. The slides were incubated in IF blocking buffer at room temperature for one hour. Primary antibodies were incubated on cells for one hour at room temperature (Supplemental methods 1C). Then cells were washed twice with PBST and incubated with secondary antibodies (Jackson Immunoresearch) for one hour at room temperature. After incubation, slides were washed with PBST three times. On the second wash, DAPI was added to the PBST. The cells were then mounted using prolong gold antifade (Molecular Probes).

### Nuclei isolation

Cells were harvested, washed twice with cold PBS, and resuspended in nuclear preparation buffer I (320 mM sucrose, 10 mM Tris (pH 8.0), 3 mM CaCl2, 2 mM Mg(OAc)2, 0.1 mM EDTA, 0.1% Triton X-100, protease inhibitors, and phosphatase inhibitors) and dounce-homogenized on ice until >95% of nuclei stained by Trypan blue. Two volumes of nuclear preparation buffer II (2.0 M sucrose, Tris (pH 8.0), 5 mM Mg(OAc)2, 5 mM DTT, 20 μM Thiamet G, protease inhibitors, and phosphatase inhibitors) were added to the nuclei suspension. Nuclei were pelleted by ultracentrifugation at 130,000 × *g* at 4°C for 45 min. Pelleted nuclei were washed with cold PBS and stored at −80°C.

### Immunoblots

Cell pellets were lysed in ice-cold RIPA buffer (150mM sodium chloride, 1% NP-40, 0.5% sodium deoxycholate, 0.1% sodium dodecyl sulfate, 50mM Tris, pH 8.0, protease inhibitors) for 20 minutes and centrifuged at 14,000 rpm for 10 minutes at 4°C. The supernatant was decanted, and Pierce 660nm Protein Assay kit (Promega) was used for quantification. Proteins were prepared using 4x Laemmli sample buffer and boiled for 5 minutes at 100°C. Proteins were separated on a denaturing SDS-PAGE gel and transferred to PVDF membrane. Membranes were blocked in PBST +5% nonfat dry milk at room temp for 30 minutes or at 4°C overnight. Membranes were incubated in primary antibodies (Supplemental methods 1C) for one hour at room temperature or at 4°C overnight and then rinsed briefly with DI water twice and washed for 5 minutes in PBST. Next, the membranes were incubated in HRP-conjugated secondary antibodies (Bio-rad) for one hour at room temperature and then washed for 5 minutes in PBST, three times. Blots were incubated with Pico Chemiluminescent Substrate (ThermoFisher) and exposed to film in a dark room.

### Sample Preparation for SILAC

LF2 (XX) and E14 (XY) mESCs were grown under standard conditions using DMEM for SILAC, 10% dialyzed FBS, 2mM glutamine, 1X non-essential amino acids, 0.1mM b-mercaptoethanol, and recombinant leukemia inhibitory factor. XX mESCs were grown in light isotopes, L-Arginine-HCL and L-Lysine-HCL. XY mESCs were grown in heavy isotopes, L-Lysine-2HCL (13C6, 15N2) and L-Arginine-HCL (13C6, 15N4) supplemented with 200 mM proline to avoid arginine-to-proline conversion. Cells were trypsinized, washed twice with cold PBS and then sonicated in 67 mM ammonium bicarbonate containing 8M guanidine HCl, 8X Phosphatase Inhibitor Cocktails II and III (Sigma-Aldrich), and 80 uM PUGNAc (Tocris Bioscience). Protein concentrations were estimated with bicinchoninic acid protein assay (ThermoFisher Scientific). Ten mgs of each lysate were combined, reduced for 1 h at 56°C with 2.55 mM TCEP and subsequently alkylated using 5 mM iodoacetamide for 45 min at room temperature in the dark. Lysates were diluted to 1M guanidine HCl using 50 mM ammonium bicarbonate, pH 8.0, and digested overnight at 37°C with sequencing grade trypsin (ThermoFisher Scientific) at an enzyme to substrate ratio of 1:50 (w/w). Tryptic peptides were acidified with formic acid (Sigma-Aldrich), desalted using a 35 cc C18 Sep-Pak SPE cartridge (Waters), and dried to completeness using a SpeedVac concentrator (Thermo).

### Lectin Weak Affinity Chromatography

Glycopeptides were enriched as described previously (Trinidad et al., 2012; Maynard et al., 2020). Briefly, desalted tryptic peptides were resuspended in 1000 μl LWAC buffer (100 mM Tris pH 7.5, 150 mM NaCl, 2 mM MgCl2, 2 mM CaCl2, 5% acetonitrile) and 100 μl was run over a 2.0 x 250-mm POROS-WGA column at 100 μl/min under isocratic conditions with LWAC buffer and eluted with a 100-μl injection of 40 mM GlcNAc. Glycopeptides were collected inline on a C18 column (Phenomenex). Enriched glycopeptides from 10 initial rounds of LWAC were eluted with 50% acetonitrile, 0.1% FA in a single 500-μl fraction, dried. LWAC enrichment was repeated for a total of three steps.

### Offline Fractionation

Glycopeptides were separated on a 1.0 × 100 mm Gemini 3μ C18 column (Phenomenex). Peptides were loaded onto the column in 20 mM NH4OCH3, pH 10 and subjected to a gradient from 1 to 21% 20 mM NH4OCH3, pH10 in 50% acetonitrile over 1.1 mL, up to 62% 20 mM NH4OCH3, pH10 in 50% acetonitrile over 5.4 mL with a flow rate of 80 μL/min.

### Mass Spectrometry Analysis

Glycopeptides were analyzed on an Orbitrap Fusion Lumos (Thermo Scientific) equipped with a NanoAcquity UPLC (Waters). Peptides were fractionated on a 15 cm × 75 μM ID 3 μM C18 EASY-Spray column using a linear gradient from 2% to 30% solvent B over 65 min. Precursor ions were measured from 350 to 1800 m/z in the Orbitrap analyzer (resolution: 120,000; AGC: 4.0e5). Each precursor ion (charged 2–8+) was isolated in the quadrupole (selection window: 1.6 m/z; dynamic exclusion window: 30 s; MIPS Peptide filter enabled) and underwent EThcD fragmentation (Maximum Injection Time: 250 ms, Supplemental Activation Collision Energy: 25%) measured in the Orbitrap (resolution: 30,000; AGC; 5.04). The scan cycle was 3 s.

Peak lists for EThcD were extracted using Proteome Discoverer 2.2. EThcD peak lists were filtered with MS-Filter, and only spectra containing a 204.0867 m/z peak corresponding to the HexNAc oxonium ion were used for database searching. EThcD data were searched against mouse and bovine entries in the SwissProt protein database downloaded on Sept 06, 2016, concatenated with a randomized sequence for each entry (a total of 22,811 sequences searched) using Protein Prospector (v5.21.1). Cleavage specificity was set as tryptic, allowing for two missed cleavages. Carbamidomethylation of Cys was set as a constant modification. The required mass accuracy was 10 ppm for precursor ions and 30 ppm for fragment ions. Variable modifications are listed in Supplemental methods 4. Unambiguous PTMs were determined using a minimum SLIP score of six, which corresponds to a 5% local false localization rate (Baker et al., 2011). Modified peptides were identified with a peptide false discovery rate of 1%. *O*-GlcNAc and *O*-GalNAc modifications were differentiated based on known protein subcellular localization and HexNAc oxonium ion fragments 138/144 ratio (Halim et al., 2014; Maynard et al., 2020).

### GFP co-immunoprecipitation

Cell pellets were lysed in ice-cold RIPA buffer supplemented with DNaseI and protease inhibitors and placed on ice for 30 minutes, extensively pipetting the suspension every 10 minutes. The suspension was centrifuged at 17,000 rpm for 10 minutes at 4°C and the supernatant lysate was decanted into a cold eppendorf tube with dilution buffer (10 mM Tris/Cl pH 7.5, 150 mM NaCl, 0.5 mM EDTA). The GFP-Trap dynabeads (Chromotek) were equilibrated with wash buffer (10 mM Tris/Cl pH 7.5, 150 mM NaCl, 0.05 % NP-40, 0.5 mM EDTA) and the lysate was bound to the beads by rotating end-over-end for one hour at 4°C. The beads were washed three times in wash buffer and bound proteins were eluted by boiling in 2x Laemmli sample buffer.

### Size Exclusion Column Chromatography

Nuclei were lysed in a high salt buffer (20mM Tris-HCl, pH 7.6, 300mM NaCl, 10% glycerol, 0.2% (v/v) Igepal (Sigma-Aldrich, 630)) and were loaded on a Superose 6 10/30GL column (Amersham) and fractionated in high salt buffer. 0.5 mL fractions were collected and concentrated into a smaller volume using a spin column-concentrator (Pierce). Fractions were then analyzed via immunoblot.

### RNA-seq

RNA was extracted using the direct-zol RNA miniprep kit (Zymo Research). Total RNA was monitored for quality control using the Agilent Bioanalyzer Nano RNA chip and Nanodrop absorbance ratios for 260/280nm and 260/230nm. Library construction was performed according to the Illumina TruSeq Total RNA stranded protocol. The input quantity for total RNA was 200ng and rRNA was depleted using ribo-zero rRNA gold removal kit (human/mouse/rat). The rRNA depleted RNA was chemically fragmented for three minutes. First strand synthesis used random primers and reverse transcriptase to make cDNA. After second strand synthesis the ds cDNA was cleaned using AMPure XP beads and the cDNA was end repaired and then the 3’ ends were adenylated. Illumina barcoded adapters were ligated on the ends and the adapter ligated fragments were enriched by nine cycles of PCR. The resulting libraries were validated by qPCR and sized by Agilent Bioanalyzer DNA high sensitivity chip. The concentrations for the libraries were normalized and multiplexed together. The multiplexed libraries were sequenced using paired end 100 cycles chemistry on the NovaSeq 6000 with a depth of 50M reads per sample.

### RNA-seq data analysis

Sequencing reads were analyzed for quality control using FASTQC(v. 0.11.2), then trimmed using Trimmomatic(v.0.32) with Illumina TruSeq adapter sequences, PHRED quality score 15 and minimum length 20 bases. The trimmed reads were aligned to the reference genome with transcriptome annotation and post-processed using Tophat2(v.2.0.12), Bowtie2(v.2.2.3), Samtools(v.0.1.19). Reads from the same samples were combined. Expression levels were quantified both with FPKM (Fragment per kilobase per million mapped reads) using Cufflinks(v. 2.1.1) and with raw counts using HTSeq(v.0.6.1p1.). Differential analysis was performed using DESeq(v.1.18.0).

### Whole Genome Bisulfite Sequencing (WGBS)

Genomic DNA was extracted using the Monarch Genomic DNA purification kit (New England Biolabs). 400ng genomic DNA was used to construct whole-genome bisulfite libraries. Briefly, DNA was sonicated by a Covaris S2 focused-ultrasonicator to ∼200bp, end repaired and adenylated using the Kapa DNA Hyper kit, and ligated to pre-methylated Illumina TruSeq LT adapters. Adapter ligated DNA was treated by the Qiagen Epitect bisulfite conversion kit. The bisulfite treated DNA was then amplified by the Bioline MyTaq HS DNA polymerase to generate final libraries for sequencing. Reads were sequenced 151×151bp paired-end on an Illumina NovaSeq 6000.

### WGBS Analysis

Raw paired-end reads were filtered for poor quality and trimmed to remove poor quality ends and adapter sequences using trim_galore (Krueger et al., 2019) v.0.5.0 with options -q 25 --stringency 3. Trimmed pairs were aligned to the mouse genome version GRCm38/mm10 using Bismark (Krueger et al., 2011) v.0.20.1 with options -L 22 -N 0 -I 0 -X 500, all other settings default. Uniquely mapping pairs were retained for analysis, and were further filtered to remove likely unconverted reads (defined as 3 consecutive unconverted Cs in the CHH context) using a custom script. PCR duplicates were removed and per-position DNA methylation information was extracted using Bismark.

DMRs were identified using the method described in Pignatta et al., 2015. Briefly, weighted average methylation at CG or non-CG sites with at least 5 coverage was obtained over non-overlapping 1kb bins tiled genome-wide for each library. Libraries were compared pairwise, for a total of 15 comparisons. For each comparison, all 1kb windows containing at least 3 informative sites shared between both samples were identified, and Fisher’s exact test was used to test for significant differences in methylation levels between the two samples. P-values were corrected for multiple hypothesis testing using the Benjamini-Hochberg (Hochberg et al., 1995) method. Windows with corrected *p* < 0.01 and a minimum difference of 35 (CG) or 10 (non-CG) percentage points were considered DMRs.

Violin plots and smoothed scatterplots were plotted in R (R Core Team., 2017) v.3.5.1 using ggplot2 (Wickham et al., 2016) v.3.1.0 and smooth Scatter respectively. Smoothed plots of DMR or feature density were obtained by counting the number of DMRs/features over 50kb windows tiled genome-wide, then smoothing using the R loess() function with span = 0.05 and plotting using ggplot2. Feature coverage was obtained by calculating the fraction of bp out of 50kb covered by the feature in each window. Smoothed plots of average methylation over chromosomes were obtained similarly by averaging methylation over 50kb windows, smoothing and plotting as described.

Major and minor satellite annotations were obtained from the UCSC table browser RepeatMasker (Smit et al., 1996) tracks for mm10, annotated as GSAT_MM and SYNREP_MM respectively. Average methylation over these regions was compared to randomly shuffled regions 200 times to obtain an approximate p-value for the significance of differences in methylation over the satellites relative to the rest of the genome. Table of alignment statistics for WGBS libraries are in Supplemental methods 5. All custom scripts are available upon request.

### RNA FISH

RNA FISH was performed using directly labeled double-stranded DNA probes, as previously described (Mlynarczyk-Evans et al., 2006). Primers used to generate the Xist probe are indicated in Supplemental methods 2A. COT-1 FISH was carried out using labeled mouse COT-1 DNA (Invitrogen). For quantification, over 100 nuclei were scored per experiment and significance was determined by using the “paired two samples for means” t-test.

### H3K27me3 IF and Xist RNA FISH combination

IF was carried out as previously described with the addition of 1mg/mL yeast tRNA (Invitrogen) in the blocking buffer and then fixed in 2% PFA for 5 minutes. Xist RNA FISH, as previously described, was immediately carried out.

### Imaging

All imaging was done on an Olympus BX60 microscope using a 100X objective. Images were collected with a Hamamatsu ORCA-ER digital camera using Micromanager software.

### Reproducibility and Rigor

All IF, immunoblots, co-IPs, and FISH are representative of at least three independent biological replicates (experiments carried out on different days with a different batch of mESCs or mEpiLCs). For targeted mESC lines, two-three independently derived lines for each genotype were assayed in at least three biological replicates. For size exclusion column chromatography assay, two biological replicates were carried out. For RNA-seq, three technical replicates of XX and three independently-derived XX-TET3KO mESCs and mEpiLCs were analyzed. For WGBS, three technical replicates of XX and three independently-derived XX-TET3KO mESCs were analyzed. We define an outlier as a result in which all the controls gave the expected outcome, but the experimental sample yielded an outcome different from other biological or technical replicates. There were no outliers or exclusions.

### Data Availability

Annotated spectra, peak lists, and the table of results for the TET1, TET2, and TET3 *O*-GlcNAcylated peptides can be viewed and downloaded from MS-Viewer with the keyword pfsmorbazl. (https://msviewer.ucsf.edu:443/prospector/cgibin/mssearch.cgi?report_title=MSViewer&search_key=pfsmorbazl&search_name=msviewer) (Baker et al., 2014).

RNA-seq data has been uploaded to GEO under accession GSE171847. WGBS data has been uploaded to GEO under accession GSE178378.

## Results

### OGT Dose Controls Cytosine Modifications

5mC and 5hmC are lower in XX mESCs than in XY mESCs when measured using assays that do not distinguish between these two modifications (Zvetcova et al., 2005; Schulz et al., 2014; Choi et al., 2017). This difference could be attributed to X-copy number, since XO mESCs exhibit XY levels of 5mC and 5hmC using these assays (Zvetcova et al., 2005; Schulz et al., 2014). Quantitation of 5mC and of 5hmC revealed a 4-fold decrease in each of these modifications in XX mESCs relative to XY mESCs (Habibi et al., 2013). To determine the effects of losing one X on 5mC and 5hmC abundance and distribution, we used immunofluorescence staining (IF) to compare these modifications in LF2 (XX), E14 (XY), and an XO line derived from LF2 (XO). 5mC (Fig. 1A) and 5hmC (Fig. 1B) levels in XY and XO cells were higher than in XX cells in serum and LIF-containing mESC media and in 2i media (Fig. 1-fig. sup. 1A-B). 5mC IF also revealed an additional difference between XX and XY/XO mESCs. In XY and XO mESCs nearly all cells showed enrichment of 5mC on pericentromeric heterochromatin, marked by intense staining with DNA dye DAPI. In contrast, in XX mESCs approximately 65% of cells showed 5mC enrichment on pericentromeric heterochromatin, while the remaining 35% of cells showed little or no pericentromeric heterochromatin enrichment (Fig.1-fig. sup. 1C). These results raised the question of why cytosine modifications in mESCs are dependent on X-copy number.

**Figure 1.**
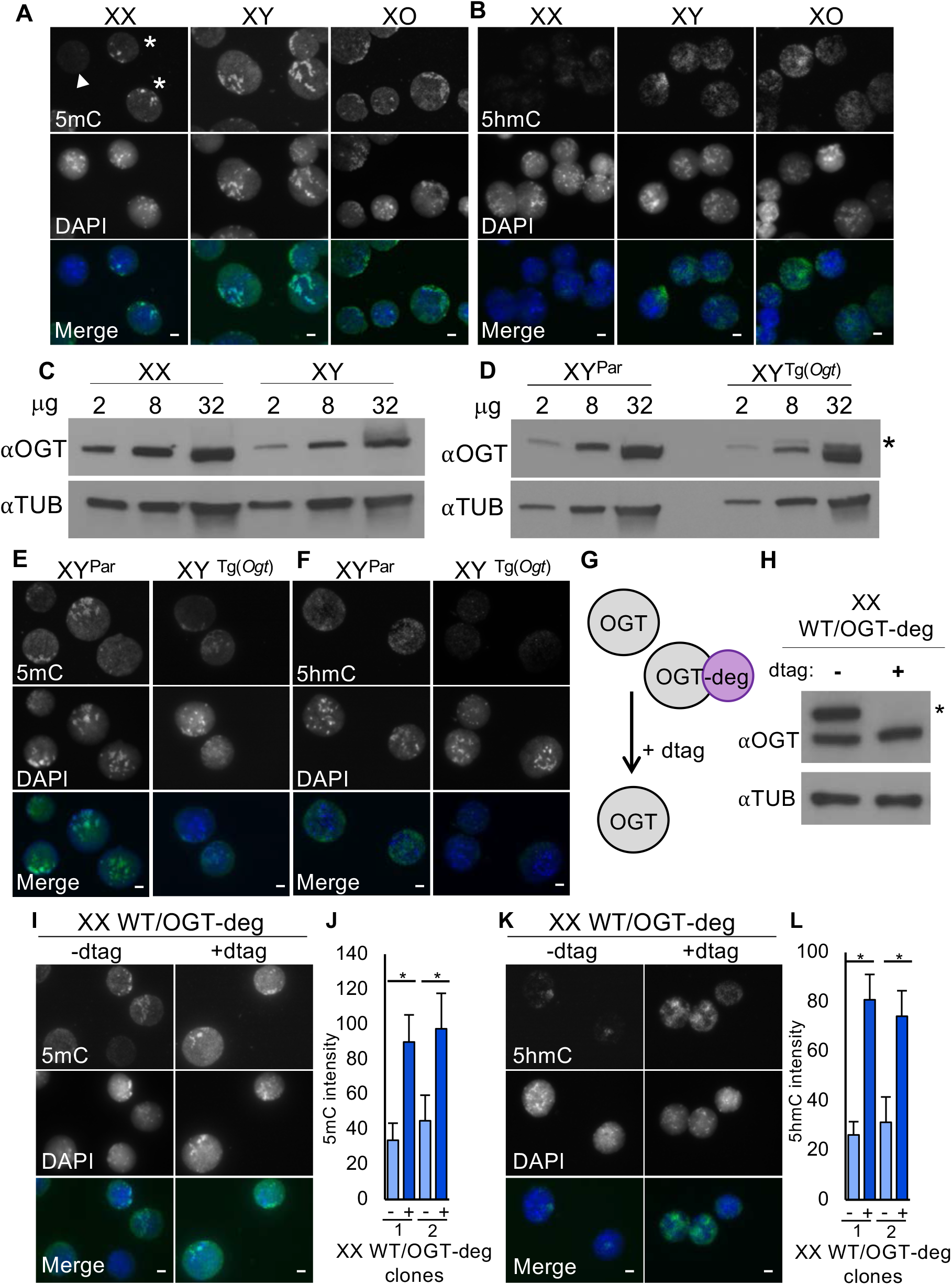
OGT dose controls cytosine modifications in mESCs. (A-B) IF staining of XX, XY, and XO mESCs using (A) 5Mc and (B) 5hmC antibodies (green in Merge). DAPI (blue in Merge) indicates nuclei. 5mC staining in XX mESCs is heterogeneous, with some cells exhibiting enrichment at pericentromeric heterochromatin (*) and some showing no apparent enrichment (arrowhead). (C-D) OGT and TUBULIN immunoblots of increasing concentration of whole cell lysate to compare OGT levels in (C) XX and XY mESCs and (D) parental XY (XY^Par^) and XY^Tg(*Ogt*)^ mESCs. * indicates transgenic FLAG-Bio tagged OGT. (E-F) IF of XY^Par^ and XY^Tg(*Ogt*)^ mESCs using (E) 5mC and (F) 5hmC antibodies (green in Merge). DAPI (blue in Merge) indicates nuclei. (G) Diagram of OGT degron (OGT-deg) strategy in XX mESCs. OGT-deg arising from the tagged *Ogt* allele is turned over by the addition of dtag. (H) OGT and TUBULIN immunoblot of XX WT/OGT-deg mESCs. * indicates degron tagged OGT. +dtag indicates a 12-hour incubation with dtag. (I and K) IF of XX WT/OGT-deg mESCs with or without 12-hour dtag incubation using (I) 5mC and (K) 5hmC antibodies (green in Merge). DAPI (blue in Merge) indicates nuclei. (J and L) Quantitation of fluorescence intensity for (I) 5mC and (K) 5hmC staining in two XX WT/OGT-deg clones treated with dtag for 0 and 12 hours, n=50 over three replicates for each condition. *=p<0.001. Scale bars in all merged images indicate 5µM.

mESCs exist in a developmental state prior to the onset of dosage compensation by XCI. Because XX mESCs have two active Xs, the increased dose of an X-linked regulator or regulators of cytosine modifications could specify the XX-specific abundance of 5mC and 5hmC. Since none of the enzymes involved in addition or turnover of 5mC are encoded on the X, an X-linked regulator of DNMTs or TETs would be a reasonable candidate. We hypothesized that OGT contributed to the XX-specific cytosine modifications since this post-translational modification enzyme is an X-linked gene product (Shafi et al., 2000) that interacts with and modifies TETs in mESCs (Vella et al., 2013; Hrit et al., 2018). OGT modification stimulates TET activity (Hrit et al., 2018) and the OGT-TET interaction has been implicated in control of TET nucleocytoplasmic distribution (Zhang et al., 2014) and chromatin association (Ito et al., 2013; Vella et al., 2013; Chen et al. 2013), providing multiple molecular mechanisms by which OGT dose could control abundance of cytosine modifications.

If OGT dose plays a role in the X-copy number dependent regulation of 5mC and 5hmC, then levels of OGT should differ between XX and XY mESCs. Immunoblot of whole cell lysates revealed that OGT was more abundant in XX than XY mESCs (Fig. 1C), consistent with its expression from two active Xs.

To determine if OGT dose affects 5mC and 5hmC levels we employed an OGT over-expressing XY mESC line (XY^Tg(Ogt)^) (Vella et al., 2013) (Fig. 1D). Using IF, we found that 5mC and 5hmC levels decrease in the XY^Tg(Ogt)^ mESC line compared to the parental XY mESC (XY^Par^) line (Fig. 1E, F). Because we were unable to generate heterozygous *Ogt* mutant XX mESCs, which is consistent with the pre-implantation lethality of *Ogt* heterozygous mutants in mice (Shafi et al., 2000), we employed a degron to change OGT dose. We introduced an inducible, FKBP degron (Nabet et al., 2018) into OGT produced from one X, generating two, independently-derived XX WT/OGT-deg clones (Fig. 1G, Fig.1-fig. sup. 1D-E), which died after five days of treatment with dtag13 (dtag) to deplete FKBP-tagged OGT. After 12 hours of treatment with dtag, FKBP-tagged OGT was no longer detectable (Fig. 1H, Fig. 1-fig. sup. 1F) and 5mC and 5hmC levels were increased (Fig. 1I-L, Fig.1-fig. sup. 1G-H). This inverse correlation between OGT levels and 5mC/5hmC abundance suggests that OGT, directly or indirectly, controls an enzyme that adds or removes cytosine DNA modifications.

### OGT differentially modifies TET3 in XX and XY mESCs

Since we investigated the role of OGT in 5mC and 5hmC accumulation because it can regulate TETs, we next asked whether TET *O*-GlcNAcylation differs in XX and XY mESCs. We employed Stable Isotope Labeling with Amino acids in Culture (SILAC) combined with lectin-weak affinity chromatography to enrich *O*-GlcNAC peptides and EThcD-MS/MS to quantitatively compare *O*-GlcNAcylated peptide abundance in XX and XY mESCs (Fig. 2A). TET3 was unique among the TETs because all of its *O*-GlcNAC peptides were upregulated in XX mESCs relative to XY mESCs (Fig. 2B-D). To determine if this increase indicated a difference in total protein, we compared TET levels in whole cell lysates of XX and XY mESCs and found that TET3, in contrast to TET1 and TET2, was more abundant in XX mESCs (Fig. 2E).

**Figure 2.**
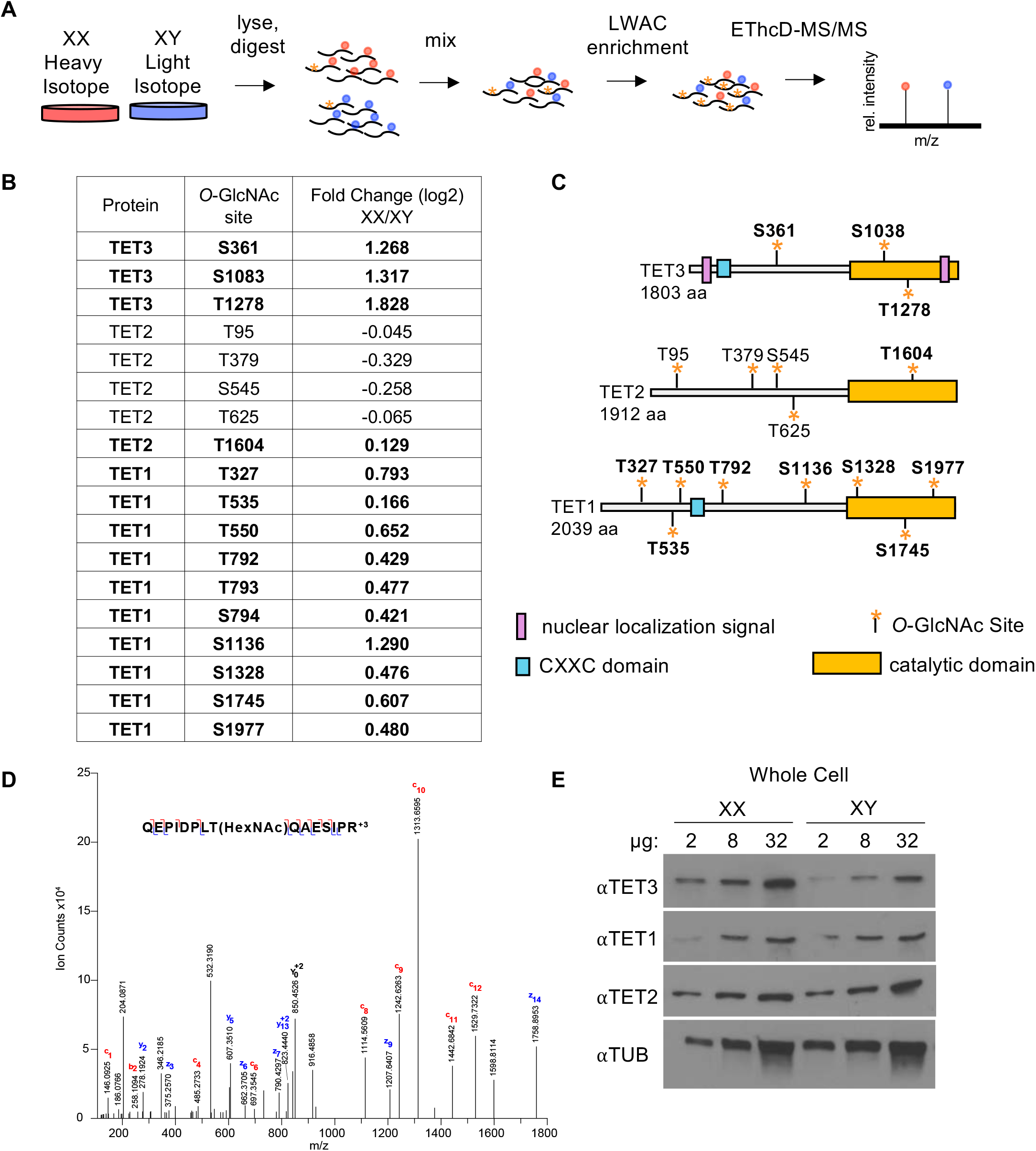
*O*-GlcNAc-modified TET3 peptides are more abundant in XX mESCs than XY mESCs. (A) Schematic of proteomic workflow: XX and XY mESCs were grown in heavy and light isotope media, respectively, and combined. Tryptic digests of combined samples were subject to lectin weak affinity chromatography (LWAC) to enrich for *O*-GlcNAcylated peptides. EThcD-MS/MS identified peptides and *O*-GlcNAc sites. (B) Table of fold change (log2) of XX/XY for each specific serine or threonine *O*-GlcNAc site found for TET1, TET2, and TET3. *O*-GlcNAc sites not previously identified (Bauer et al., 2015) are bolded. (C) Diagram of mouse TET proteins, yellow star indicates *O*-GlcNAc site, blue box indicates CXXC domain, yellow box indicates catalytic domain, purple box indicates TET3 nuclear localization signal, KKRK (Xiao et al., 2013). (D) The spectrum of TET3 *O*-GlcNAcylated at T1278. (E) TET1, TET2, TET3, and TUBULIN (TUB) immunoblots of increasing concentration of whole cell lysate prepared from XX and XY mESCs.

### OGT and TET3 accumulate in the nucleus of XX mESCs but not in XY mESCs

OGT can control TET3 nucleocytoplasmic distribution (Zhang et al., 2014) prompting us to investigate TET3 distribution in XX and XY mESCs. IF revealed that TET3 was predominantly cytoplasmic in XY mESCs and predominantly nuclear in XX mESCs (Fig. 3A). This nuclear accumulation occurred in an additional XX mESC line (PGK12.1) and was not dependent on growth media, as it was observed in mESC media and 2i media (Fig. 3-fig. sup. 1A-B). Immunoblotting for TET3 in nuclear extracts showed that more TET3 was present in XX nuclei than XY nuclei (Fig. 3B), consistent with the IF results. Similarly, OGT was predominantly localized to the nucleus in XX and to the cytoplasm in XY mESCs (Fig. 3C-D).

**Figure 3.**
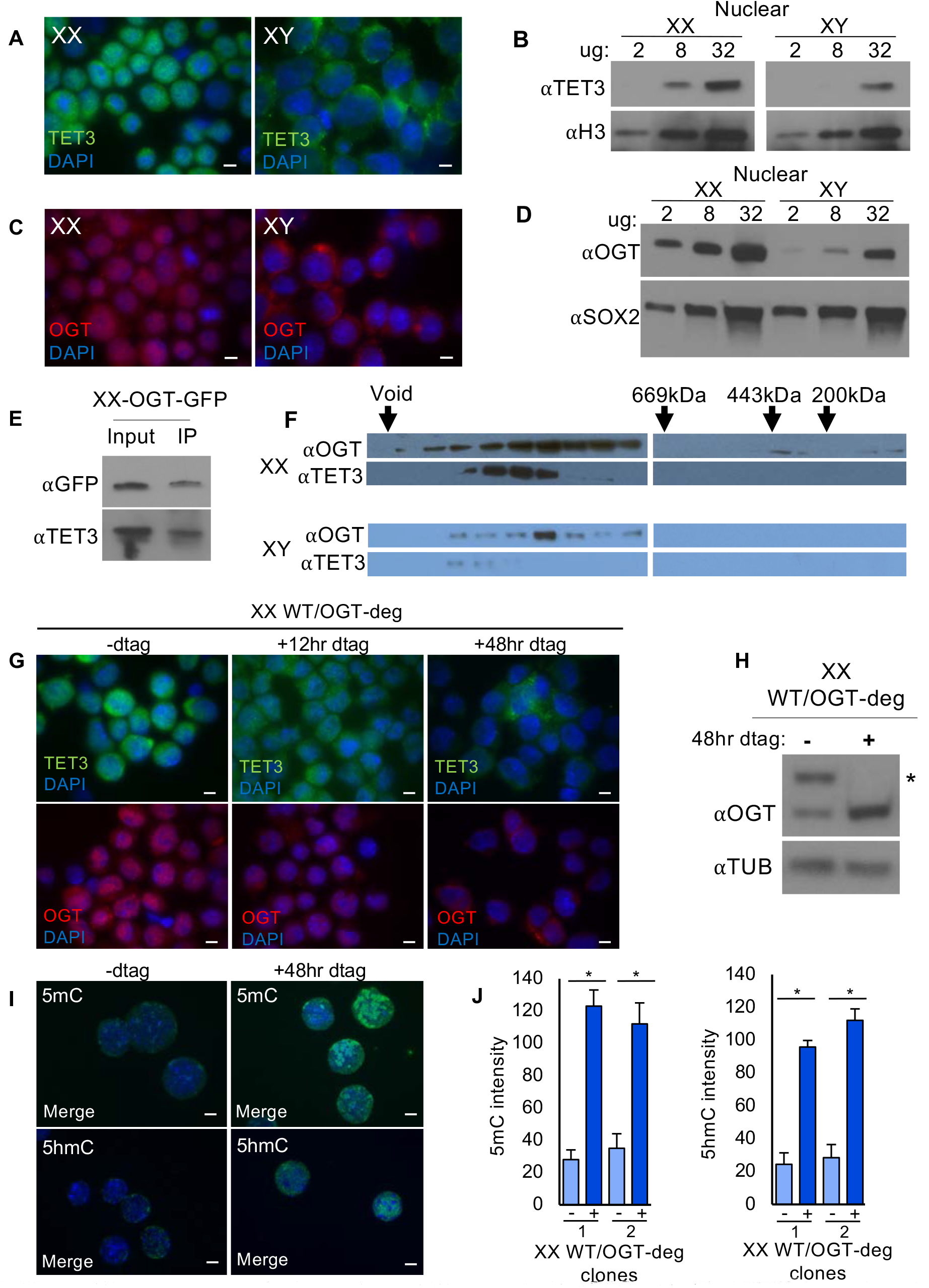
OGT and TET3 accumulate in the nucleus of XX but not XY mESCs. (A) IF of TET3 (green) in XX and XY mESCs. DAPI (blue) indicates nuclei. (B) TET3 and histone H3 (H3) immunoblots of increasing concentration of nuclear lysate prepared from XX and XY mESCs. (C) IF of OGT (red) in XX and XY mESCs. DAPI (blue) indicates nuclei. (D) OGT and SOX2 immunoblots of increasing concentration of nuclear lysate prepared from XX and XY mESCs. (E) OGT tagged with GFP (endogenously on both Xs); XX-OGT-GFP was immunoprecipitated with a GFP monoclonal antibody. Immunoblots show input and IP probed with TET3 and GFP monoclonal antibody. (F) OGT and TET3 immunoblots of fractions from size exclusion chromatography of nuclear extracts prepared from XX and XY mESCs. Size of molecular weight markers indicated by arrows. (G) IF of TET3 (green) and OGT (red) in (G) XX WT/OGT-deg mESCs with or without 12hr and 48hr dtag incubation. DAPI (blue) indicates nuclei. (H) OGT and TUBULIN immunoblot of XX WT/OGT-deg mESCs. * indicates degron tagged OGT. +dtag indicates 48-hour incubation. (I) IF of XX WT/OGT-deg mESCs with or without 48-hour dtag incubation using 5mC and 5hmC antibodies (green in Merge). DAPI (blue in Merge) indicates nuclei. (J) Quantitation of fluorescence intensity for 5mC and 5hmC staining in two XX WT/OGT-deg clones treated with or without dtag for 48 hours, n=50 over three replicates for each condition. *=p<0.001. Scale bars in all merged images indicate 5µM.

OGT and TET3 interact when co-expressed in transformed human cells (Ito et al., 2014). We queried whether XX mESCs also showed this interaction using co-immunoprecipitation (co-IP) and co-fractionation. For co-IP experiments we generated an XX mESC line in which both alleles of *Ogt* were tagged with GFP (Fig 3.-fig. sup. 1C-D). TET3 associated with OGT-GFP in IPs generated with a GFP antibody (Fig. 3E). Size exclusion column chromatography of nuclear extracts showed that OGT and TET3 fractionate together in high molecular weight complexes in both XX and XY mESCs (Fig. 3F). However, the fractions containing OGT and TET3 differed between XX and XY mESC nuclear extracts, suggesting that the OGT-TET3 complex composition is affected by the number of Xs and/or the presence of the Y chromosome.

In addition to OGT, XX, and XY mESCs have the potential to differ in dose of the majority of X-linked genes, any number of which could control TET3 nucleocytoplasmic distribution. To determine whether OGT dose affects TET3 localization we used a time course to assess the effects of OGT turnover on TET3 distribution in XX WT/OGT-deg mESCs. After 12 hours of dtag incubation, TET3 no longer appeared distinctly nuclear. After 48 hours it was observed predominantly in the cytoplasm and there was a decrease in TET3 abundance (Fig. 3G). Like TET3, OGT also appeared to be predominantly cytoplasmic after 48 hours in dtag (Fig. 3G). While dtag-treated XX WT/OGT-deg mESCs exhibited lower levels of OGT by IF, immunoblot showed that after 48 hours of turnover of the OGT-deg the expression of wildtype OGT increased (Fig. 3H, Fig. 3-fig. sup. 1E), potentially reflecting differences in epitope availability in IF and immunoblot. As was observed for 12 hours of dtag treatment, 5mC and 5hmC levels were increased after 48 hours in dtag (Fig. 3I-J). The decrease in TET3 observed after 48 hours of dtag treatment may contribute to this increase in abundance of modified cytosines. In XY^Tg(Ogt)^ mESCs the increased OGT dose correlated with increased abundance of TET3 and OGT in the nucleus (Fig. 3- fig. sup. 1F). Together these results indicate that the higher OGT dose in XX mESCs promotes nuclear accumulation of TET3 and OGT.

### XX mESCs lacking TET3 exhibit increased 5mC and 5hmC levels

We reasoned that XX-specific nuclear enrichment of TET3 may provide the basis for lower levels of 5mC and 5hmC in XX mESCs. To test this hypothesis, we mutated both copies of *Tet3* in XX mESCs (Fig. 4-fig.sup 1-3) generating XX-TET3KO mESCs. The XX-TET3KO mESCs exhibited altered colony morphology, growing in tight colonies that expanded more in the vertical than horizontal direction (Fig 4-fig. sup. 4A). Because mutation of TET1 in XX mESCs changed the abundance of TET2 (Hrit et al., 2018), we first examined whether homozygous mutation of *Tet3* affected steady state levels of the other TETs. In XX-TET3KO mESCs TET3 and TET1 were undetectable, while TET2 levels were not appreciably altered (Fig. 4A), indicating that TET3 is necessary for TET1 accumulation in XX mESCs. 5mC and 5hmC levels in XX-TET3KO mESCs were elevated relative to XX mESCs and enrichment of 5mC at pericentromeric heterochromatin was not as apparent in XX-TET3KO mESCs (Fig. 4B-E). Upon quantitation of fluorescent intensity, 5mC and 5hmC levels in all three XX-TET3KO clones were significantly greater (p<0.001) than XX mESCs (Fig. 4C and E).

**Figure 4.**
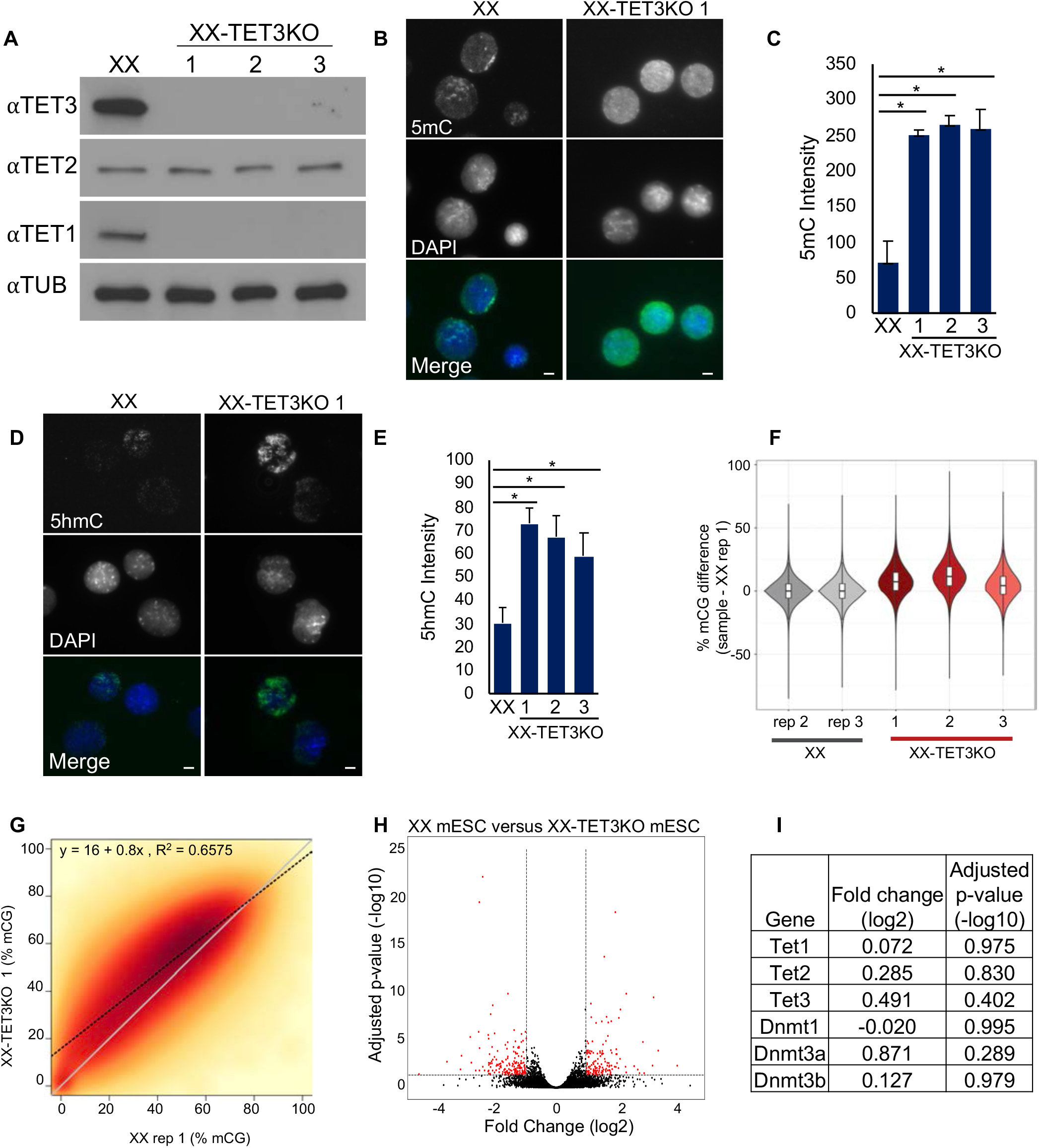
Loss of TET3 alters cytosine modifications without significantly changing gene expression in XX mESCs. (A) TET1-3 and TUBULIN (TUB) immunoblots of whole cell lysate from wildtype (XX) and 3 independently-derived TET3 knock-out (XX-TET3KO) XX mESCs. TET3 antibody is directed to a C-terminal epitope which is downstream of the guide cut sites that lie in the first coding exon. (B and D) IF of (B) 5mC (green in Merge) and (D) 5hmC (green in Merge) in XX and XX-TET3KO 1 mESCs. DAPI (blue in Merge) indicates nuclei. Scale bars in all merged images indicate 5µm. (C and E) Fluorescence quantitation of (B) 5mC and (D) 5hmC staining in XX and XX-TET3KO clones. * indicates statistically significant difference, p<0.001. Error bars indicate sd, n=50 over three replicates for each condition. (F) Difference in WGBS average CG methylation percentage between one XX replicate (rep) and the remaining samples, over 1kb regions tiled genome-wide. (G) Smoothed scatterplot of average methylation over 1kb bins tiling the entire genome in XX (x-axis) vs. XX-TET3KO 1 (y-axis). Line of best fit indicated by black dotted line, with corresponding equation and R^2^ value at top of each plot. y=x line plotted in grey for reference. (H) Volcano plot of RNA-seq data from XX versus XX-TET3KO clones showing genes with fold changes (log2) >-4 and <4. 3 genes are excluded. Red dots indicate genes with fold changes (log2) < -1 or > 1 and adjusted p-value <0.01. (I) Table of fold change (log2) and adjusted p-value (-log10) of cytosine modification enzymes in XX versus XX-TET3KO mESC clones.

To obtain a high-resolution view of the effects of *Tet3* mutation, we performed whole genome bisulfite sequencing (WGBS) which simultaneously detects 5mC and 5hmC. In all three XX-TET3KO mESC clones, 5mC+5hmC was globally increased compared to three replicates of XX mESCs (Fig. 4F-G, Fig.4- fig. sup. 5A-B). Differentially modified region (DMR) analysis over 1kb non-overlapping windows tiled genome-wide identified 6,741 DMRs, which almost exclusively had more 5mC+5hmC in XX-TET3KOs than XX mESCs (Fig. 4- fig. sup. 6A-B). When comparing the X to autosomes, we observed that DMRs were slightly but consistently depleted from the X and that the X was less modified overall, in both XX and XX-TET3KO mESCs (Fig. 4- fig. sup. 7). While 5mC+5hmC were increased on the X in XX-TET3KOs, the increase was less than the increase on autosomes. Analysis of major and minor satellite repeats showed that unlike major satellites in the rest of the genome, minor satellites did not appreciably increase levels of 5mC+5hmC (Fig. 4- fig. sup. 8A-B), consistent with the uniform 5mC staining throughout the nucleus in XX-TET3KO mESCs. Together, these bisulfite-seq and IF results indicate that TET3 and TET1 are necessary to maintain correct levels of 5mC+5hmC and correct distribution of 5mC in XX mESCs.

To ask whether the changes in cytosine modifications observed in XX-TET3KO mESCs impacted gene expression, we performed RNA-seq. 104 genes were up regulated in the XX-TET3KO mESCs relative to XX controls, and 86 were down regulated (Fig. 4H, Fig. 4- fig. sup. 4B). Cytosine modification pathway transcripts were not significantly changed in XX-TET3KO mESCs (Fig. 4I). Gene Ontology (GO) term analysis of the 190 genes with altered expression showed no statistically significant results.

### XX-TET3KO mEpiLCs exhibit altered Xist RNA distribution

XX and XY mESCs are pluripotent and have the potential to differentiate into all cell types of the developing and adult mouse embryo (Reik et al., 2015). To ask whether the XX-specific nuclear enrichment of TET3 is necessary for a developmental transition, we differentiated XX and XX-TET3KO mESCs into mEpiLCs. Both cell types exhibited comparable morphology at day 5 of differentiation (Fig.5- fig. sup.1A). While the differentiating XX-TET3KO mESCs initially grew normally, after day 4 they exhibited more cell death than their wildtype counterparts. Comparison of XX and XX-TET3KO mEpiLC RNA-seq at day 5 of differentiation showed that 404 genes exhibited increased expression and 499 exhibited decreased expression in XX-TET3KO mEpiLCs (Fig. 5A, Fig. 5-fig. sup. 1B). The expression of mESC markers (naive pluripotency genes) went down and mEpiLC markers (formative and/or primed pluripotency genes) went up comparably upon differentiation of XX and XX-TET3KO mESCs (Fig. 5B), suggesting that many key transcriptional changes that characterize this transition can occur when TET3 is knocked out. Despite up-regulation of mEpiLC markers, GO term analysis showed gene expression changes affecting growth and development (Fig. 5C), suggesting an altered developmental program. Upon differentiation, IF showed that 5mC and 5hmC in XX-TET3KO mEpiLCs were comparable to XX mEpiLCs (Fig 5-fig. sup. 1C-D), consistent with similar changes in steady state RNA levels of *Dnmt*s and *Tet*s in both genotypes (Fig. 5-fig. sup. 1E).

**Figure 5.**
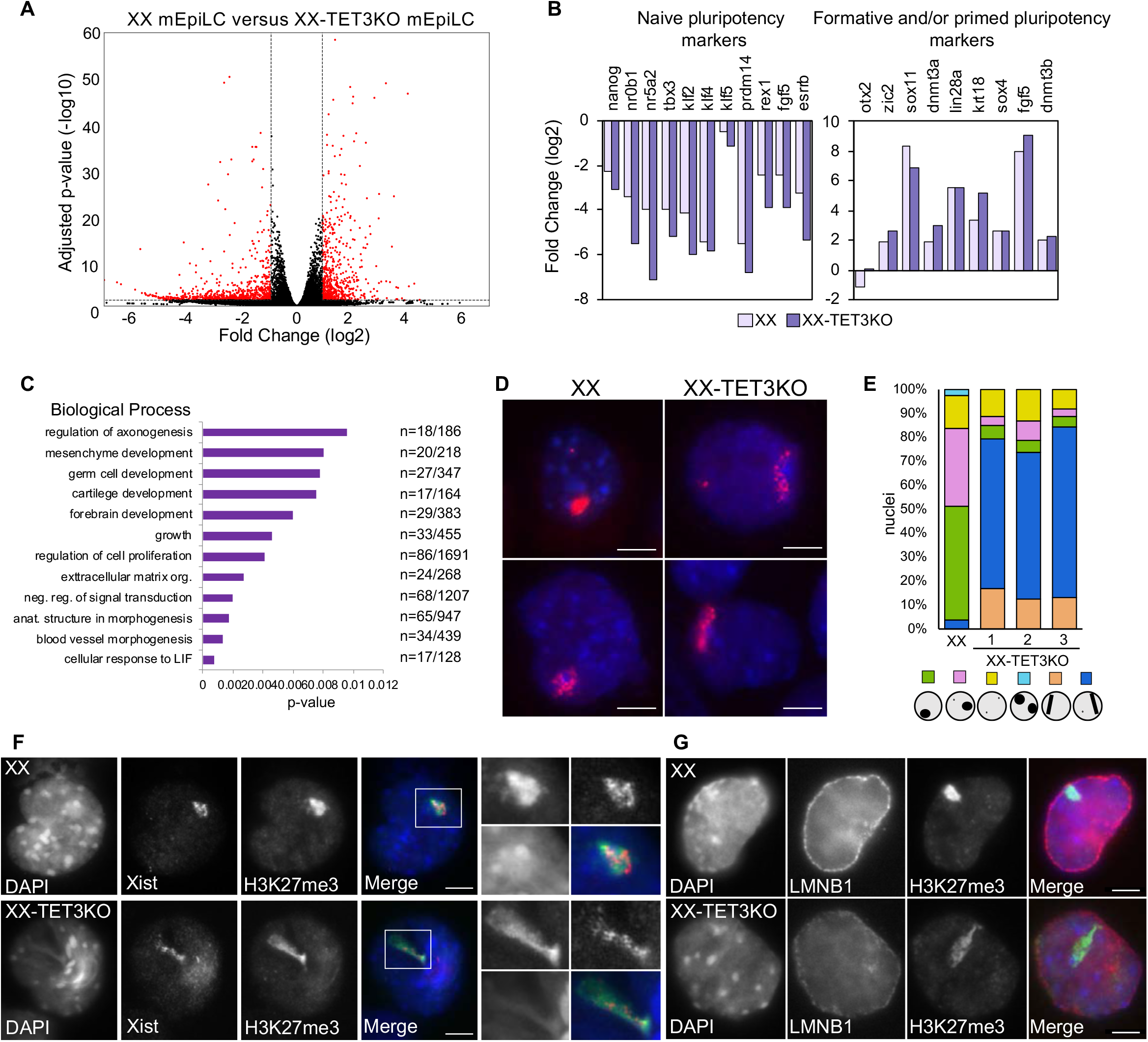
Loss of TET3 alters gene expression and Xist RNA distribution in mEpiLCs. (A) Volcano plot of RNA-seq data from XX versus XX-TET3KO mEpiLC clones showing genes with fold changes (log2) >-7 and <7 and adjusted p-value (-log10) >60. Red dots indicate genes with fold changes (log2) < -1 or > 1 and adjusted p-value (-log10) <0.01. (B) Fold change (log2) between day 0 and day 5 of differentiation for selected naïve and formative/primed pluripotency markers in XX (light purple) and XX-TET3KO (dark purple) cells. (C) GO terms (biological process) enriched for genes differentially expressed between XX and XX-TET3KO mEpiLCs. Only GO terms with FDR p-values <0.01 are included. n indicates the number of differentially expressed genes in the data set/the number of genes that define that GO term. (D) The predominant pattern of Xist RNA FISH (red) in XX and XX-TET3KO 1 at day 5 of mEpiLC differentiation. DAPI (blue) indicates nuclei. (E) Proportion of cells exhibiting different Xist RNA distribution patterns in XX and XX-TET3KO 1 mEpiLCs, n>100 cells/replicate. (F) Xist RNA FISH (red in Merge) combined with IF for H3K27me3 (green in Merge) in XX and XX-TET3KO 1 mEpiLCs. DAPI (blue in Merge) indicates nuclei. The white box indicates the location of the zoom-in for each image. For quantitation see Fig. 5- fig. sup. 1F. (G) IF of H3K27me3 (green in Merge) and LAMIN B1 (LMNB1; red in Merge) in XX and XX-TET3KO 1 mEpiLCs. DAPI (blue in Merge) indicates nuclei. For quantitation see Fig. 5- fig. sup. 1G. White scale bar in all merged images indicates 5µm.

The mESC to mEpiLC transition is similar to the transition that occurs upon implantation *in vivo*, and is accompanied by significant epigenetic changes (Takahashi et al., 2018). One of these changes is XCI, which occurs in XX, but not XY cells. The onset of XCI is marked by the up-regulation and *cis*-spread of the X-linked Xist non-coding RNA from its site of transcription to cover, or coat, the X that will be silenced (Boeren et al., 2021). *Xist* is silenced by 5mC/5hmC in differentiating XY mESCs, since the aberrant up-regulation and *cis*-spread of Xist RNA occurs in differentiating XY mESCs that have dramatically reduced 5mC/5hmC due to loss of DNMT1 (Panning et al., 1996). This result indicates that the elevated 5mC and/or 5hmC in XY mESCs inhibits the up-regulation and *cis*-spread of Xist RNA and raises the possibility that the increase in cytosine modifications in XX-TET3KO mESCs may impact Xist RNA. To examine this possibility, we performed Xist RNA fluorescence *in situ* hybridization (FISH) in XX and XX-TET3KO mEpiLCs. In XX mEpiLCs, Xist RNA was present in puncta packed into spherical clouds typical of coating in the majority of cells (Fig 5. D-E). In XX-TET3KO mEpiLCs Xist RNA distribution was markedly different, showing a more dispersed pattern, appearing as a linear or comet-like cloud in the majority of cells (Fig 5. D-E). These results show that TET3 is necessary for normal Xist RNA distribution at the onset of XCI.

Xist RNA recruits the Polycomb Repressive Complex 2, which results in an increase in H3K27 tri-methylation (H3K27me3) on the Xist RNA coated X (Sarma et al., 2014). To ask whether loss of TET3 alters the ability of Xist RNA to promote H3K27me3 we performed Xist RNA FISH together with IF for this mark in XX and XX-TET3KO mEpiLCs (Fig. 5F). The Xist RNA and H3K27me3 signals overlapped in the majority of cells in both XX (97%) and XX-TET3KO (91%) mEpiLCs (Fig. 5-fig. sup. 1F), indicating that TET3 is not necessary for Xist RNA to recruit H3K27me3 enzymes.

In differentiated XX cells, Xist RNA is often observed at the nuclear periphery, in contact with the nuclear lamina (Rego et al., 2008). To ask whether the aberrantly distributed Xist RNA in XX-TET3KO mEpiLCs exhibits association with the nuclear lamina, we performed IF for H3K27me3 and the nuclear lamina marker LAMIN B1 (LMNB1). In both XX mEpiLCs and in XX-TET3KO the region of H3K27me3 enrichment was localized to the LMNB1-stained nuclear lamina (Fig. 5G) in greater than 85% of the cells (Fig. 5-fig. sup. 1G). In XX-TET3KO mEpiLCs, LMNB1 levels appeared to be lower than in XX cells. LMNB1 transcripts were comparable between genotypes, suggesting any differences on the protein level cannot be attributed to steady state RNA differences (Fig. 5-fig. sup. 1H). These H3K27me3 and LMNB1 data suggest that despite its aberrant distribution, Xist RNA in XX-TET3KO mEpiLCs is able to mediate changes in chromatin structure and nuclear organization that are characteristic of silencing.

To ask whether Xist RNA was able to cause other chromosome-wide alterations associated with silencing, we examined whether exclusion of RNA polymerase II (pol II) or histone H3 lysine 4 di-methylation (H3K4me2) occurred in XX-TET3KO mEpiLCs. While pol II and H3K4me2 were excluded from the H3K27me3-enriched region in XX mEpiLCs, such exclusion was rarely seen in the XX-TET3KO mEpiLCs (Fig. 6A-D). To determine whether the comet-like Xist RNA coating was able to silence genes, we examined the distribution of COT-1 containing transcripts (Fig. 6E-F). COT-1 consists of repetitive sequences that are present in introns and thus excluded from the inactive X (Hall et al., 2010). FISH for Xist and COT-1 RNA showed that COT-1 containing transcripts were excluded from the Xist RNA cloud in the majority of XX mEpiLCs. In contrast, COT-1 containing transcripts were coincident with Xist RNA in the majority of XX-TET3KO mEpiLCs. Consistent with the silencing defect, a larger proportion of X-linked genes than autosomal genes in the XX-TET3KO mEpiLCs exhibited a 2-fold or greater increase in steady-state RNA levels when compared to XX mEpiLCs (Fig. 6G). This analysis of RNA-seq data, in combination with the data showing that the transcriptional machinery, a chromatin mark associated with transcriptional activity, and nascent transcripts are no longer robustly excluded from the Xist RNA or H3K27me3-enriched domain in *Tet3* mutants, indicate that the aberrantly distributed Xist RNA in XX-TET3KO mEpiLCs does not efficiently silence X-linked genes.

**Figure 6.**
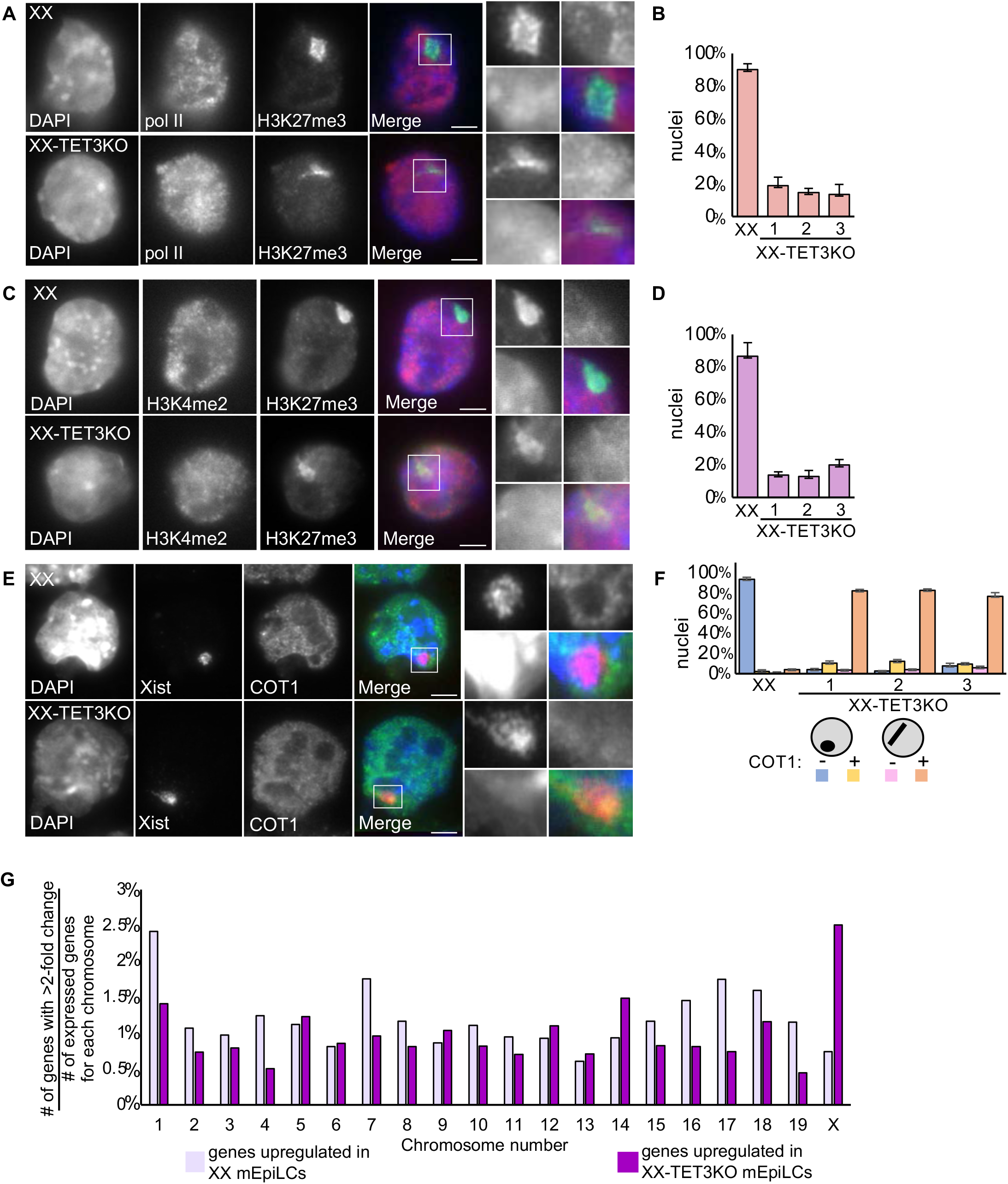
Xist RNA function is disrupted in XX-TET3KO mEpiLCs. (A) IF of RNA polymerase II (pol II; red in Merge) and H3K27me3 (green in Merge) in XX and XX-TET3KO 1 mEpiLCs. DAPI (blue in Merge) indicates nuclei. The white box indicates the location of the zoom-in for each image. (B) Proportion of nuclei exhibiting pol II exclusion for XX and XX-TET3KO clones. Error bars indicate sd, n>100 cells/replicate. (C) IF of H3K4me2 (red in Merge) and H3K27me3 (green in Merge) in XX and XX-TET3KO 1 mEpiLCs. DAPI (blue in Merge) indicates nuclei. (D) Proportion of cells exhibiting H3K4me2 exclusion in XX and XX-TET3KO clones. Error bars indicate sd, n>100 cells/replicate. (E) Xist RNA FISH (red in Merge) combined with COT1 RNA FISH (green in Merge) in XX and XX-TET3KO 1 mEpiLCs. DAPI (blue in Merge) indicates nuclei. (F) Proportion of cells exhibiting exclusion of COT1 RNA in Xist RNA-enriched regions. Error bars indicate sd, n>100/replicate. White scale bars in all merged images indicate 5µm. (G) The proportion of genes with a fold change (log2) of <-1 (dark purple) or > 1 (light purple), padj (log -10) < 0.01 in XX mEpiLCs vs XX-TET3KO mEpiLCs, normalized to the total number of expressed genes per chromosome.

## Discussion

### OGT regulates levels of modified cytosines in mESCs

Our results show that OGT dose affects levels of 5mC and 5hmC in mESCs. Specifically, increased expression of OGT in XY mESCs resulted in a decrease in these cytosine modifications, while decreased expression in XX mESCs resulted in an increase in both these modifications. These results implicate OGT as one of the X-linked genes that contributes to the decreased levels of 5mC and 5hmC observed in XX mESCs relative to XY/XO mESCs. The phosphatase DUSP9 is another X-linked regulator of cytosine modifications, since heterozygous deletion of *Dusp9* in XX mESCs increases 5mC levels (Choi et al., 2017). Our data raise the possibility that OGT may be one of the factors that DUSP9 signaling influences to regulate cytosine modifications.

Since OGT activity is regulated by levels of its cofactor UDP-GlcNAc, the inverse relationship between OGT dose and abundance of cytosine modifications provides a potential connection between the cell’s metabolic state and the epigenome. These results raise the interesting possibility that inputs into UDP-GlcNAc production, like glucose and glutamine, may impact gene expression by controlling cytosine modification states. In addition, differences in access to glucose and glutamine within colonies may contribute to the variability in intensity and distribution of 5mC staining in XX mESCs.

### OGT-modified TET3 is more abundant in XX than XY mESCs

While OGT has thousands of potential targets, our unbiased mass spectrometry approach identified TET3 as a cytosine modification enzyme that is differentially modified by OGT in XX vs XY mESCs. The higher abundance of *O*-GlcNAcylated TET3 peptides in XX mESCs reflected an increase in the total amount of TET3. In addition, TET3 in XX mESCs was largely nuclear, while it was largely cytoplasmic in XY mESCs. The lower abundance and predominantly cytoplasmic localization of TET3 in XY mESCs is consistent with near-undetectable expression of endogenously tagged TET3 in this cell type (Pantier et al., 2019) and may explain why TET3 makes a smaller contribution to total 5hmC than TET1 and TET2 in XY mESCs (Dawalty et al., 2014). Since *de novo* methyltransferases are more abundant in XY mESCs (Zvetcova et al., 2005; Schulz et al., 2014) (Fig 7- fig. sup. 1A) and TET3 resides in the nucleus and is more highly expressed in XX mESCs, the X-copy number dependent differences in 5mC/5hmC likely reflect the differing balance between addition and turnover activities in these pluripotent stem cells. Because OGT modifies many additional histone post-translational modification enzymes and chromatin remodelers (Myers et al., 2011), it is possible that the epigenetic differences between XX and XY mESCs are not restricted to cytosine modifications.

### OGT dose determines intracellular distribution of TET3 and OGT

The XX-specific nuclear accumulation of TET3 and OGT depends on the XX dose of OGT, since turnover of OGT arising from one X results in the redistribution of TET3 and OGT to the cytoplasm (Fig. 7A). TET3 may be preferentially localized in the nucleus due to its increased *O*-GlcNAcylation in XX mESCs. This post-translational modification could be acting on the sequences necessary for nuclear export or import. The dose-dependent nuclear enrichment of OGT in XX mESCs may be dependent on its association with TET3 or on its *O*-GlcNAcylation (Kreppel et al., 1997). In contrast to this finding in mESCs, OGT inhibits nuclear accumulation of TET3 when these proteins are co-transfected into HeLa cells (Zhang et al., 2014), raising the possibility that TET3 distribution is controlled in a cell-type specific manner. OGT interacts with TET3 and these proteins occur in differing high molecular weight complexes in XX and XY mESCs. Determining what other proteins are in the complexes may give insight as to why OGT and TET3 are localized to the nucleus in XX mESCs.

**Figure 7.**
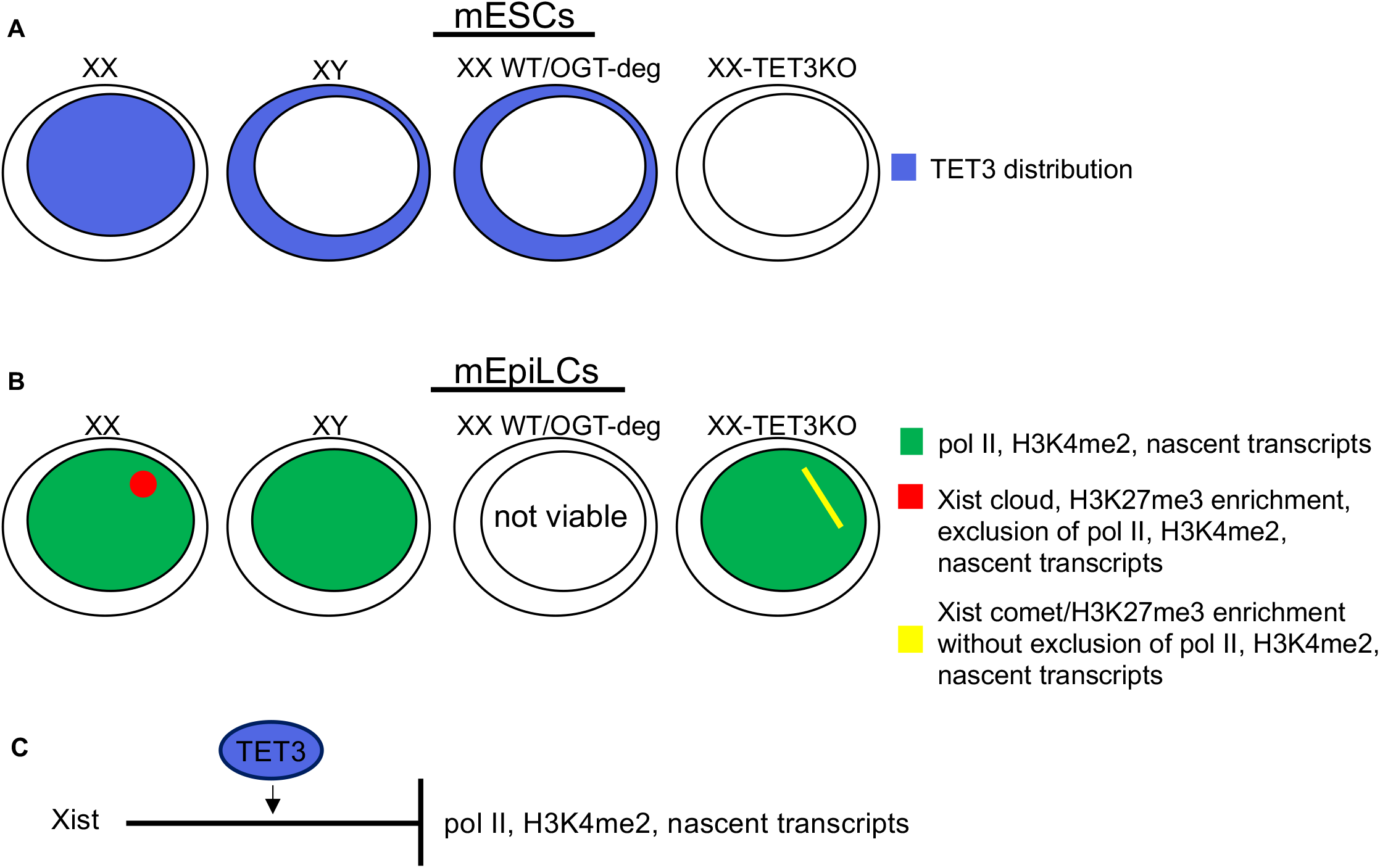
TET3 exhibits X-dose dependent distribution in mESCs and is necessary for Xist RNA-mediated silencing upon differentiation. (A) TET3 is enriched in the nucleus in XX mESCs and enriched in cytoplasm of mESCs that express OGT from one X (XY and XX WT/OGT-deg mESCs). (B) Xist RNA is distributed in a spherical, cloud-like shape and H3K27me3 is enriched in the same region (red) while pol II, H3K4me2, and nascent transcripts are excluded from that region (green) in wildtype XX mEpiLCs. In the XX-TET3KO mEpiLC clones, Xist RNA is distributed in a linear, comet-like shape, which is coincident with H3K27me3 enrichment, from which pol II, H3K4me2, and nascent transcripts are not excluded. Yellow indicates the overlap between Xist RNA/H3K27me3 (red) and pol II/H3K4me2/nascent transcripts (green). (C) TET3 is necessary Xist RNA mediated silencing, as measured by exclusion of pol II, H3K4me2, and nascent transcripts.

### Mutation of *Tet3* affects 5mC/5hmC abundance in XX mESCs

Knockout of TET3 in XX mESCs resulted in loss of TET1 and increased abundance of 5mC and 5hmC without notable changes in gene expression. The distribution of these marks was similar between XX and XX-TET3KO mESCs, with an overall increase in the non-repetitive regions of the mutant genomes. The X exhibited lower modifications than autosomes in both genotypes, though the difference was less marked in the XX-TET3KOs. Allele-specific analysis would be necessary to determine if one or both Xs is the source of the overall reduction in 5mC+5hmC relative to autosomes. In addition the heterogeneity of 5mC staining was reduced and enrichment of 5mC at pericentromeric heterochromatin was less marked in XX-TET3KO mESCs. Because both TET3 and TET1 are no longer appreciably detectable in the XX-TET3KO mESCs, it is impossible to distinguish which enzyme to attribute particular changes in 5mC and 5hmC to. TET1 steady state mRNA levels are not appreciably altered in XX-TET3KO mESCs, suggesting post-transcriptional regulation of TET1 accumulation. In XX mESCs, a mutation in TET1 that impairs its interaction with OGT results in increased abundance of TET2 without a change in *Tet2* mRNA levels (Hrit et al., 2018). Understanding how the abundance of all three TETs is connected, and if this connection is unique to pluripotent cells with two Xs, may provide useful insights into TET biology.

### XX-TET3KO mEpiLCs exhibit changes in Xist RNA distribution and silencing

In XX and XX-TET3KO mEpiLCS naive pluripotency markers were decreased and formative/primed pluripotency markers were increased, indicating that both genotypes achieved key hallmarks of differentiation into mEpiLCs. Despite similar changes in pluripotency marker expression, XX and XX-TET3KO mEpiLCs exhibited expression differences in growth and developmental pathways, consistent developmental failure observed upon germline deletion of TET3, TET1, or all three TETs (Gu et al., 2011; Dai et al., 2016; Khoueiry et al., 2017).

While achieving hallmarks of mEpiLC differentiation, distribution and activity of Xist RNA was altered in XX-TET3KO mEpiLCs. A spherical Xist RNA cloud was predominantly seen in wildtype XX mEpiLCs, while an elongated comet-like cloud was seen most often in XX-TET3KO mEpiLCs. Despite this unusual distribution, H3K27me3 was enriched where Xist RNA localizes and the region of H3K27me3-enrichment, presumably reflecting the Xist coated X, associates with the nuclear lamina. In wildtype mEpiLCs, regions of H3K27me3 enrichment or Xist RNA accumulation are associated with silencing, as shown by exclusion of pol II, H3K4me2, and nascent transcripts. In contrast, these three indicators of transcriptional activity are not excluded in XX-TET3KO mEpiLCs (Fig. 7B-C).

XCI is coupled to differentiation, since normal differentiation of XX mESCs to mEpiLCs requires XCI (Schulz et al., 2014). In addition, Xist RNA can coat but not silence the X in XY mESCs expressing an inducible Xist cDNA transgene if expression is induced after 2 days of embryoid body or retinoic acid-induced differentiation (Wutz et al. 2000). Together, these findings suggest that there is a precise developmental window during which the X is silencing competent, and that a silencing-induced signal in XX mESCs is necessary for further differentiation. Perhaps the changes in the proteome or epigenome associated with differentiation occur with different kinetics when XX-TET3KO mESCs are directed to the mEpiLC fate, affecting one or both of these processes.

Mutational analysis of *Xist* revealed that Initial establishment of silencing during XCI involves at least two steps: the formation of a compartment that contains Xist RNA and that is devoid of transcriptional machinery, followed by the shift of transcriptionally active genes into this silencing compartment. (Wutz et al., 2002; Chaumeil et al., 2006). A repetitive region of Xist, the E-repeat, is necessary for formation of the Xist RNA-containing silencing compartment. The E-repeat nucleates a protein condensate that is critical for silencing, by virtue of its interaction with the RNA binding protein CELF1 and associated proteins (Pandya-Jones et al., 2020). Xist RNA in mEpiLCs derived from *Tet3* mutant mESCs exhibits similarities to E-repeat mutant Xist RNA, which also shows an altered distribution and silencing defects (Pandya-Jones et al., 2020). The similarity between the *Tet3* mutant and *Xist E-repeat* mutant phenotypes may indicate that both perturbations affect the same pathway. While expression of proteins associated with Xist RNA mediated silencing is not significantly altered in XX-TET3KO mESCs (Fig. 7- fig. sup. 1B), perhaps a protein that is necessary for CELF1 complex activity is aberrantly expressed or exhibits altered function in differentiating XX-TET3KO mESCs. Alternatively, epigenetic changes to the underlying chromatin may make the X in differentiating XX-TET3KO mESCs an unsuitable substrate for silencing.

### Does the OGT-TET3 complex ensure silencing occurs only in XX cells?

In flies and worms, which also equalize X-linked gene dosage between the sexes, the dosage compensation pathway is controlled by complexes that are sensitive to the ratio of X to autosomal components (Cline et al., 1996). The activities of these complexes differ when the X:autosome ratio is 1:1 (XX) or 0.5:1 (XY in flies or XO in worms). As a result, a two-fold difference in X-linked components can be converted into an all-or-none response and the expression of the factors that mediate dosage compensation depends on X dosage. Our data suggest that the OGT-TET3 complex may participate in mammalian X-dosage sensing, since it differs in distribution and composition in XX and XY mESCs and since TET3 is necessary for Xist RNA-mediated silencing.

## Acknowledgements

We thank the Vella Lab for the XY^Par^ and XY^Tg(Ogt)^ mESC lines. We thank the Genomics High Throughput Facility Shared Resource of the Cancer Center Support Grant (P30CA-062203) at the University of California, Irvine and NIH shared instrumentation grants 1S10RR025496-01, 1S10OD010794-01, and 1S10OD021718-01 for RNA-seq library preparation, sequencing, and data analysis. We thank Mahnaz Akhavan and the Broad Stem Cell Research Center BioSequencing Core for high throughput sequencing. We thank all members of the Panning lab, Elphege Nora, Vijay Ramani, and Kate Carbone for valuable ideas and discussion. This work was supported by R01 GM128431-02 (BP), F32 GM136115 (CLP), the Sandler Program for Breakthrough Biomedical Research (BP) (AB), the Dr Miriam & Sheldon G. Adelson Medical Research Foundation (AB), the Howard Hughes Medical Institute (AB) (SEJ).

---

**Figure 1- figure supplement 1.**
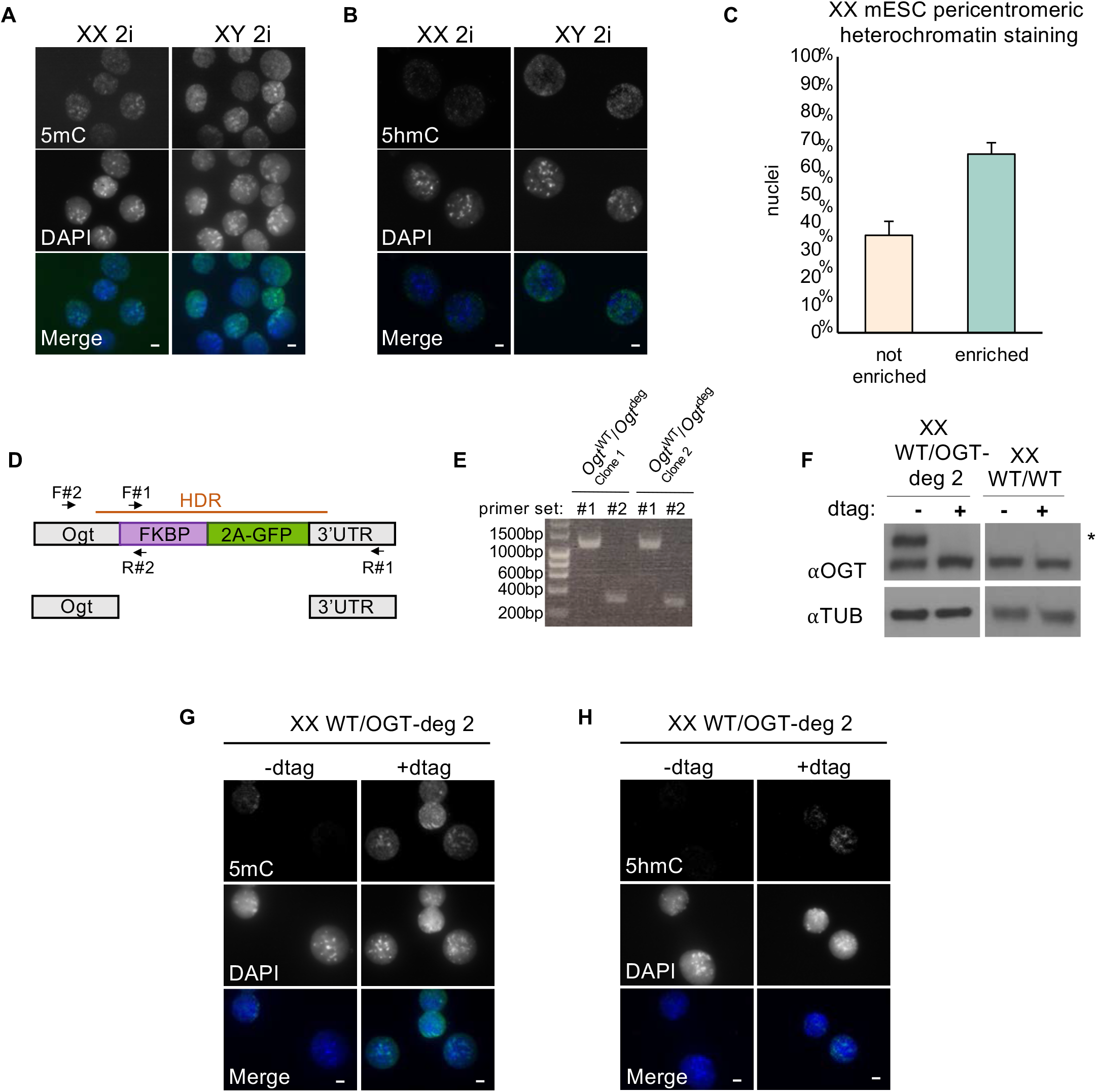
(A-B) IF of XX and XY mESCs grown in 2i using (B) 5mC and (C) 5hmC antibodies (green in Merge). DAPI (blue in Merge) indicates nuclei. (C) Proportion of XX mESC cells exhibiting enriched pericentromeric heterochromatin staining of 5mC, n>100. Error bars indicate sd. (D) Schematic of XX WT/OGT-deg genotype. DNA encoding the FKBP degron (deg) was added to the 3’ end of one *Ogt* allele, followed by a 2A sequence and green fluorescent protein sequence (GFP). The 2A sequence causes ribosome skipping, resulting in separate translation of OGT-deg and 2A-GFP. Orange line: template used for homology-directed repair (HDR). Horizontal arrows: primers used for PCR genotyping. (E) PCR genotyping of two independently derived, and targeted XX mESC clones using primers indicated in (D). (F) OGT and TUBULIN (TUB) immunoblot of second, independently-derived clone of XX WT/OGT-deg and XX WT/WT mESCs. In WT/OGT-deg cells the lower band represents OGT arising from the wild type allele and the larger band (*) is consistent with the anticipated 13 kDA increase introduced by the degron tag. +dtag indicates a 12-hour incubation with dtag. (G and H) IF of second, independently-derived XX WT/OGT-deg clone with or without 12-hour dtag incubation using (G) 5mC and (H) 5hmC antibodies (green in Merge). DAPI (blue in Merge) indicates nuclei. White scale bars in all merged images indicate 5µm.

**Figure 3- figure supplement 1.**
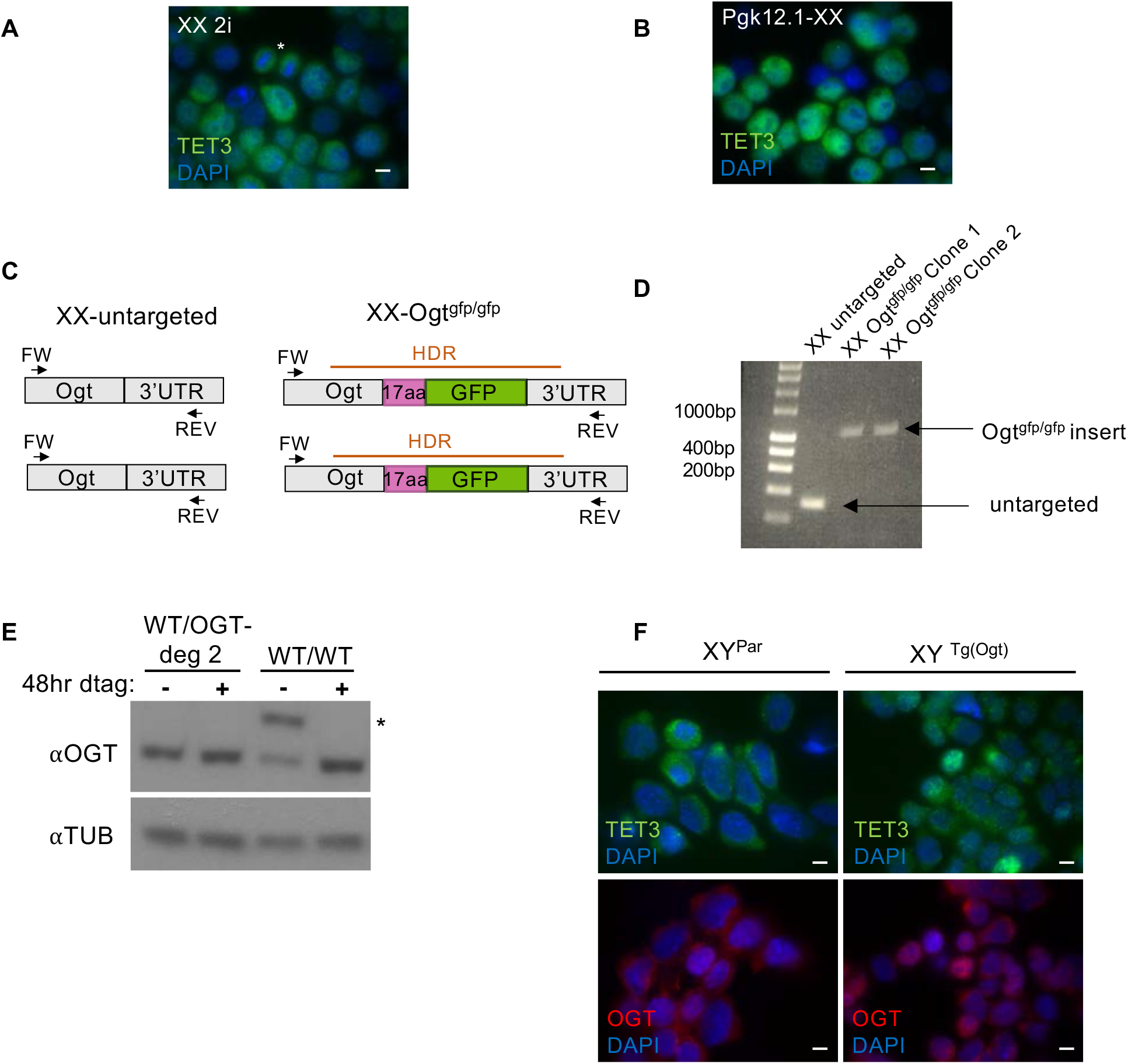
(A) IF for TET3 (green) in XX mESCs grown in 2i media. DAPI (blue) indicates nuclei, White star indicates mESC in mitosis. (B) IF of TET3 (green) in PGK12.1, a second XX mESC line. DAPI (blue) indicates nuclei. (C) Schematic of XX-*Ogt^gfp/gfp^* genotype. DNA encoding a 17 amino acid flexible linker and green fluorescent protein (GFP) was added to the 3‘end of both *Ogt* alleles. Orange line: template used for homology-directed repair (HDR). Horizontal arrows: primers used for PCR genotyping. (D) PCR genotyping of two independently derived, and targeted mESC clones using primers indicated in (C). (E) OGT and TUBULIN (TUB) immunoblot of XX WT/WT mESCs and a second independently-derived clone of XX WT/OGT-deg. In WT/OGT-deg cells the lower band represents OGT arising from the wild type allele and the larger band (*) is consistent with the anticipated 13 kDA increase introduced by the degron tag. +dtag indicates a 48-hour incubation with dtag. (F) IF of TET3 (green) and OGT (red) in XY^Par^ and XY^Tg(Ogt)^ mESCs. DAPI (blue) indicates nuclei. Scale bars in all merged images indicate 5µm.

**Figure 4- figure supplement 1.**
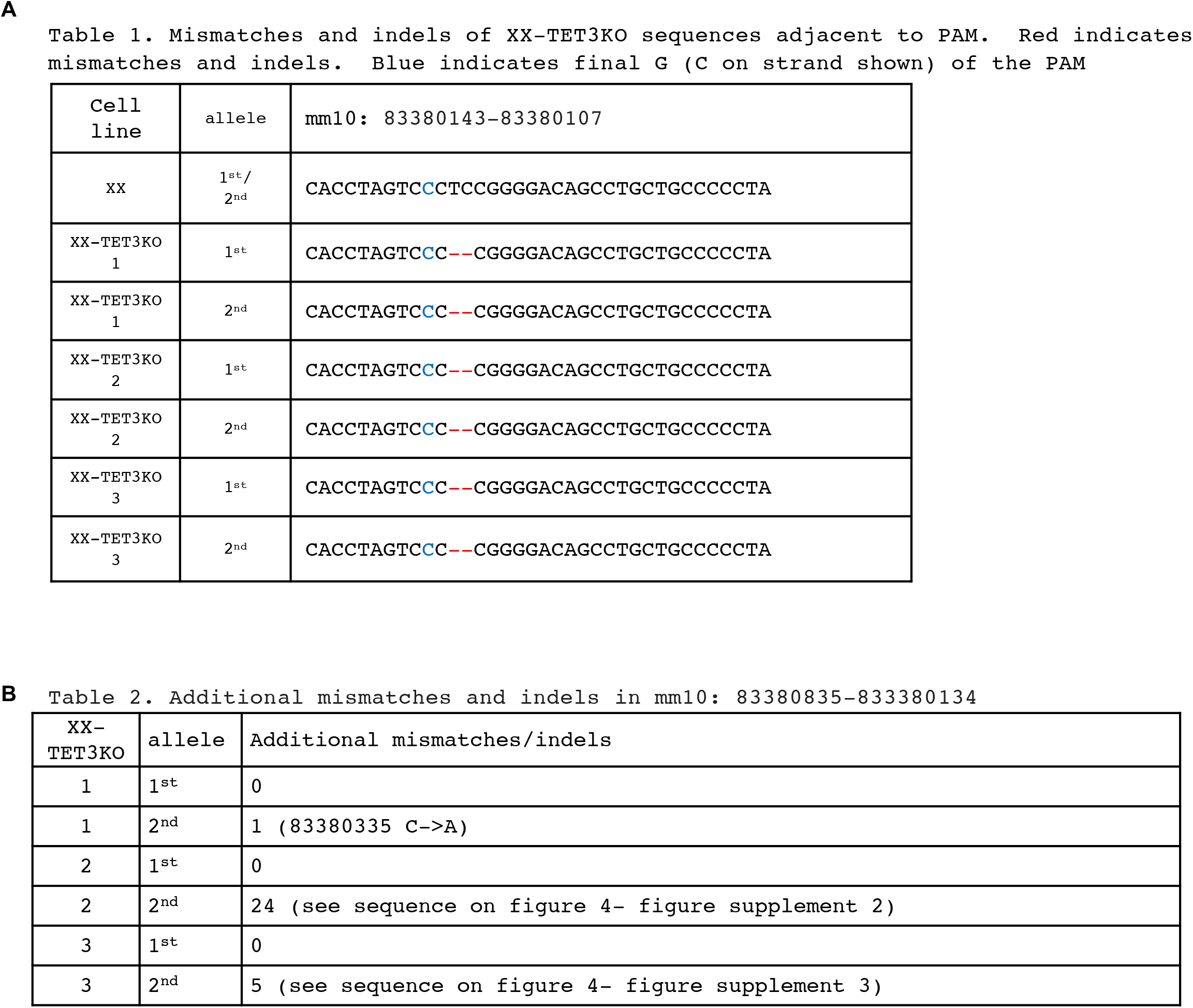
(A) Table 1 of mismatches and indels of XX-TET3KO sequences adjacent to PAM. Red font indicates mismatches and indels. Blue font indicates final G (C on strand shown) of PAM. Mm10 refers to the NCBI GRC38 *Mus. musculus* genome. (B) Table 2 of additional mismatches and indels within the PAM regions.

**Figure 4- figure supplement 2.**
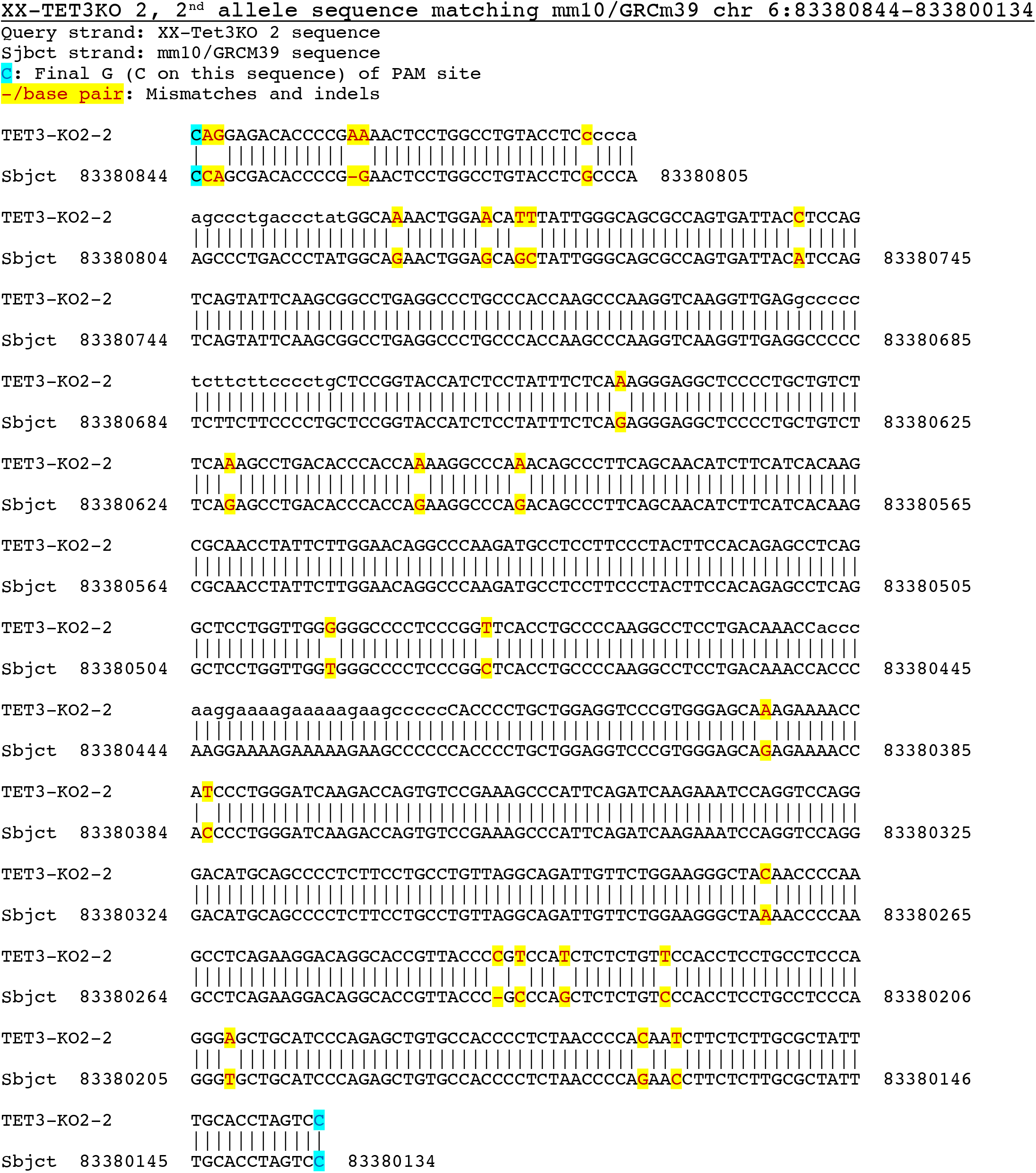
Aligned sequences of XX-TET3KO alleles to mm10/GRCm39 genome. Query strand indicates XX-TET3KO PCR product using FW and REV primers 40 base pairs up or downstream of the PAM site. Sbjct strand indicates the mm10/GRCm39 *mus musculus* genome. Blue text and highlight indicates the final G (C on strand) of the PAM site. Red text and yellow highlight indicate mismatches and indels between the PAM sites.

**Figure 4- figure supplement 3.**
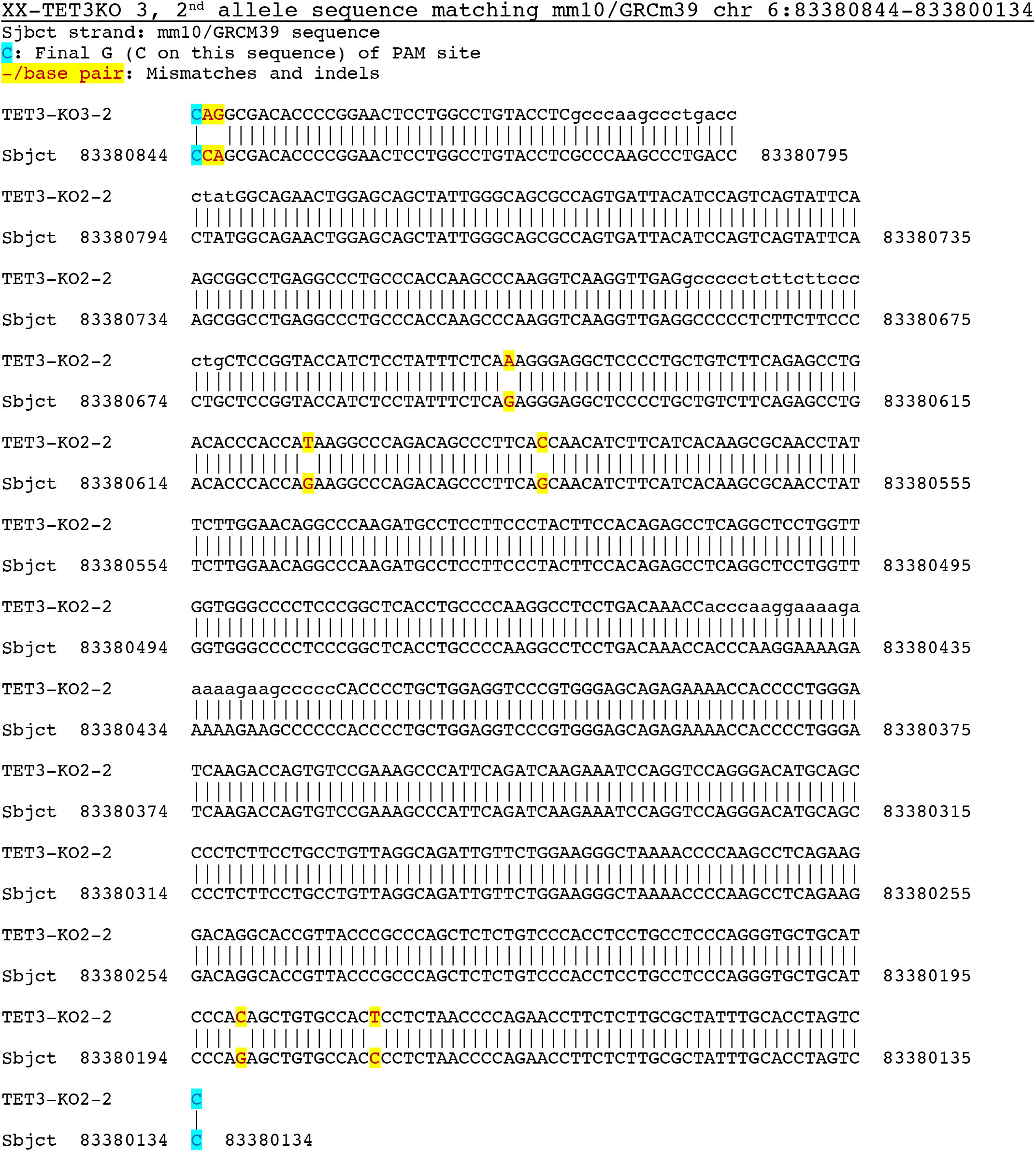
Aligned sequences of XX-TET3KO alleles to mm10/GRCm39 genome. Query strand indicates XX-TET3KO PCR product using FW and REV primers 40 base pairs up or downstream of the PAM site. Sbjct strand indicates the mm10/GRCm39 *mus musculus* genome. Blue text and highlight indicates the final G (C on strand) of the PAM site. Red text and yellow highlight indicate mismatches and indels between the PAM sites.

**Figure 4- figure supplement 4.**
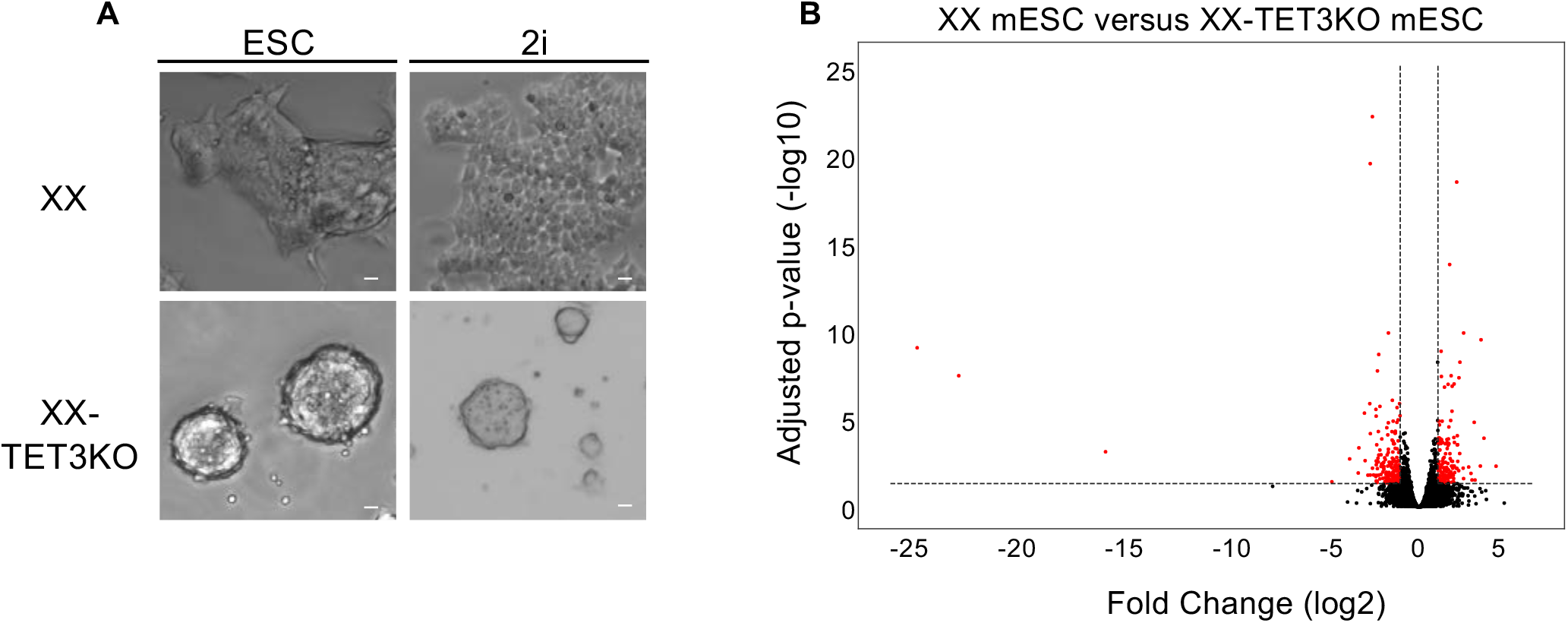
(A) Brightfield images of XX and XX-TET3KO mESCs grown in ESC or 2i media. In contrast to the normal colony morphology of XX mESCs, XX-TET3KO mESCs exhibit towering colonies which grow vertically regardless of media type. White scale bars indicates 5µM. (B) Full volcano plot comparing RNA-seq from XX and XX-TET3KO mESCs showing genes excluded in Fig. 4H. Red dots indicate genes with fold changes (log2) < -1 or > 1 and adjusted p-value <0.01.

**Figure 4- figure supplement 5.**
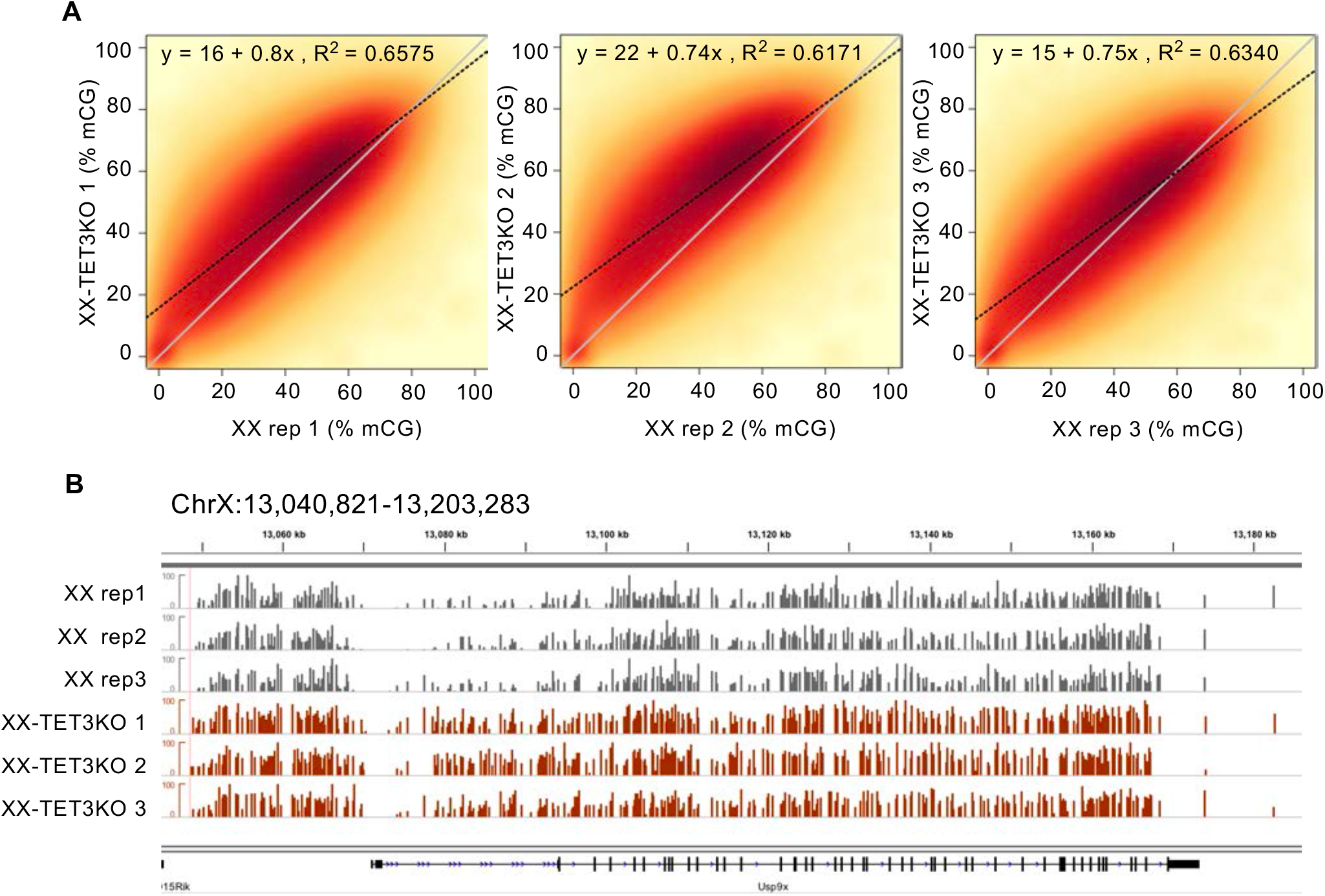
WGBS of XX-TET3KO versus XX mESCs. (A) Smoothed scatterplot of average methylation over 1kb bins tiling the entire genome in XX (x-axis) vs. XX-TET3KO (y-axis). Line of best fit indicated by black dotted line, with equation and R^2^ value at top of each plot. y=x line also plotted in grey for reference. Scale from 0- 100 (%CG methylation) (B) Example region on ChrX showing hypermethylation in XX-TET3KO lines relative to XX.

**Figure 4- figure supplement 6.**
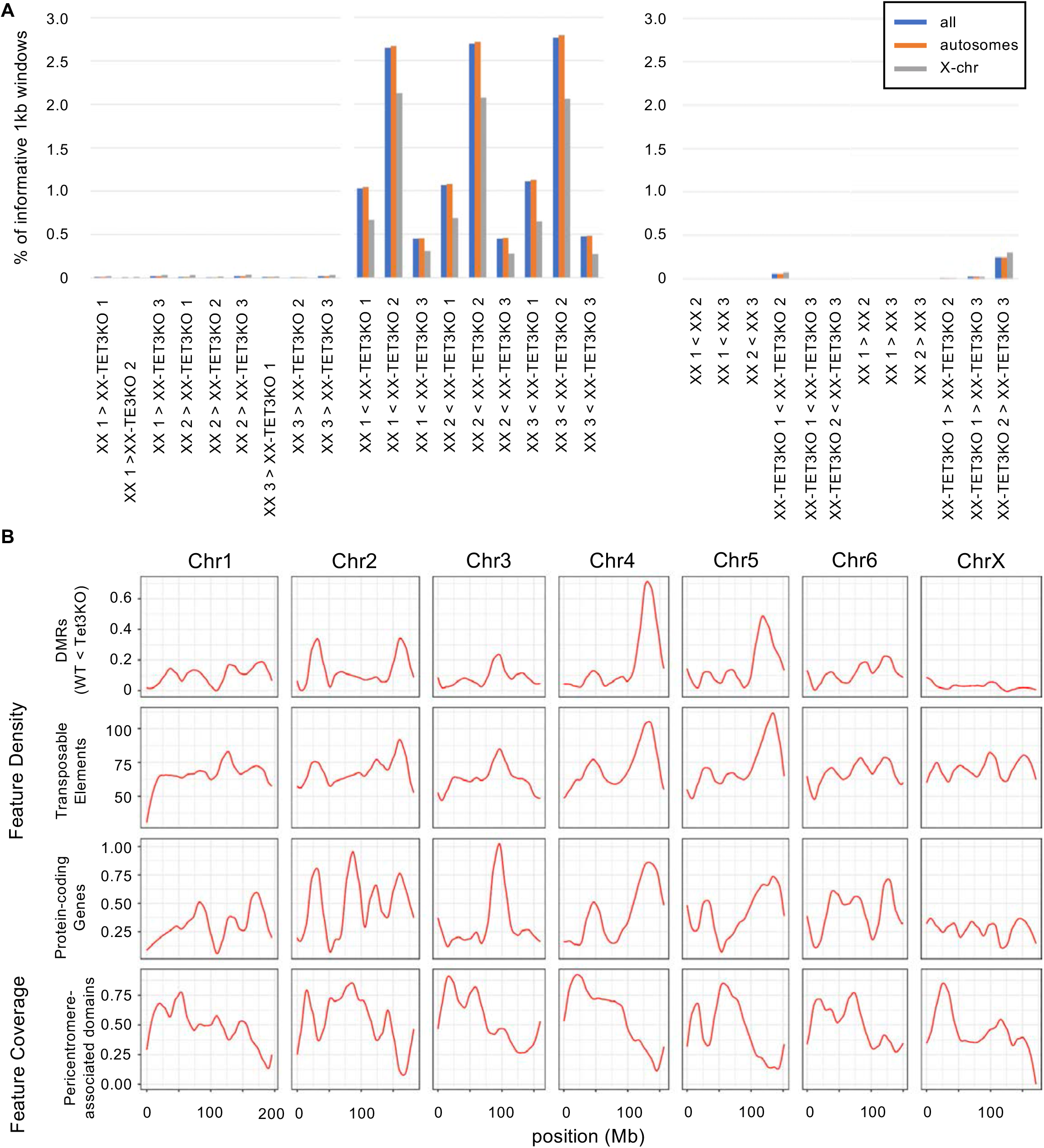
DMR analysis of XX-TET3KO WGBS data. (A) Percent of all informative 1kb windows tiled genome-wide that were identified as significant DMRs for each set of comparisons. Plots on left show XX vs. XX-TET3KO, plots on right show within-genotype (false positive) DMRs. (B) Distribution of the 6,741 ‘consensus’ XX < XX-TET3KO DMRs (identified as DMRs in at least 5/9 pairwise comparisons between all XX and XX-TET3KO replicates), relative to other common genomic features: transposons, protein-coding genes, and pericentromere associated domains (Wijchers et al. 2015). Plots show loess-smoothed average feature density (number of features) or feature coverage (average % of bp covered by feature) for 50kb windows tiled genome-wide. First six autosomes + X shown.

**Figure 4- figure supplement 7.**
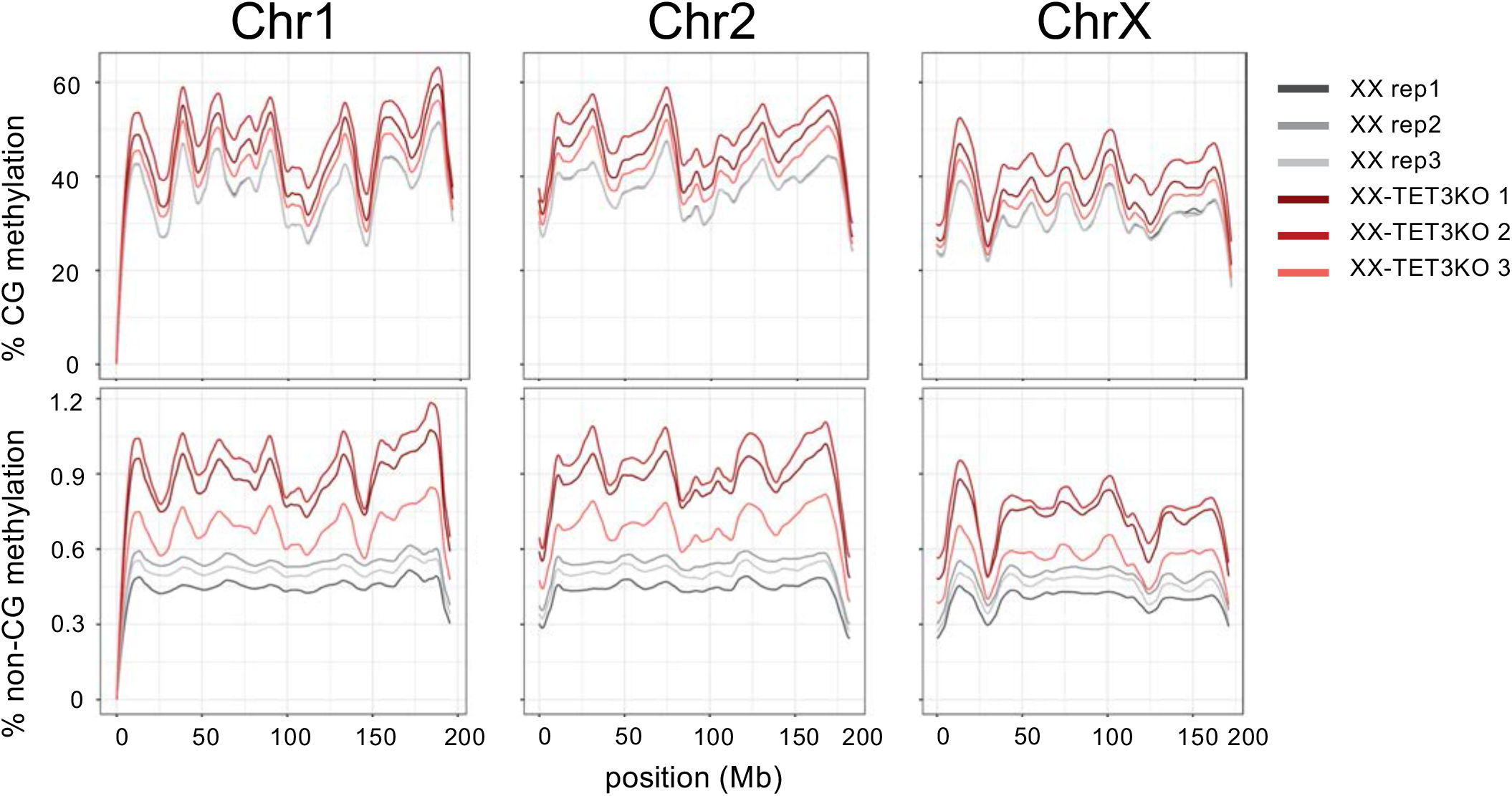
Loess-smoothed average CG (top) and non-CG (bottom) methylation (5mC + 5hmC) over chromosomes 1, 2, and X. Methylation was averaged over 50kb windows.

**Figure 4- figure supplement 8.**
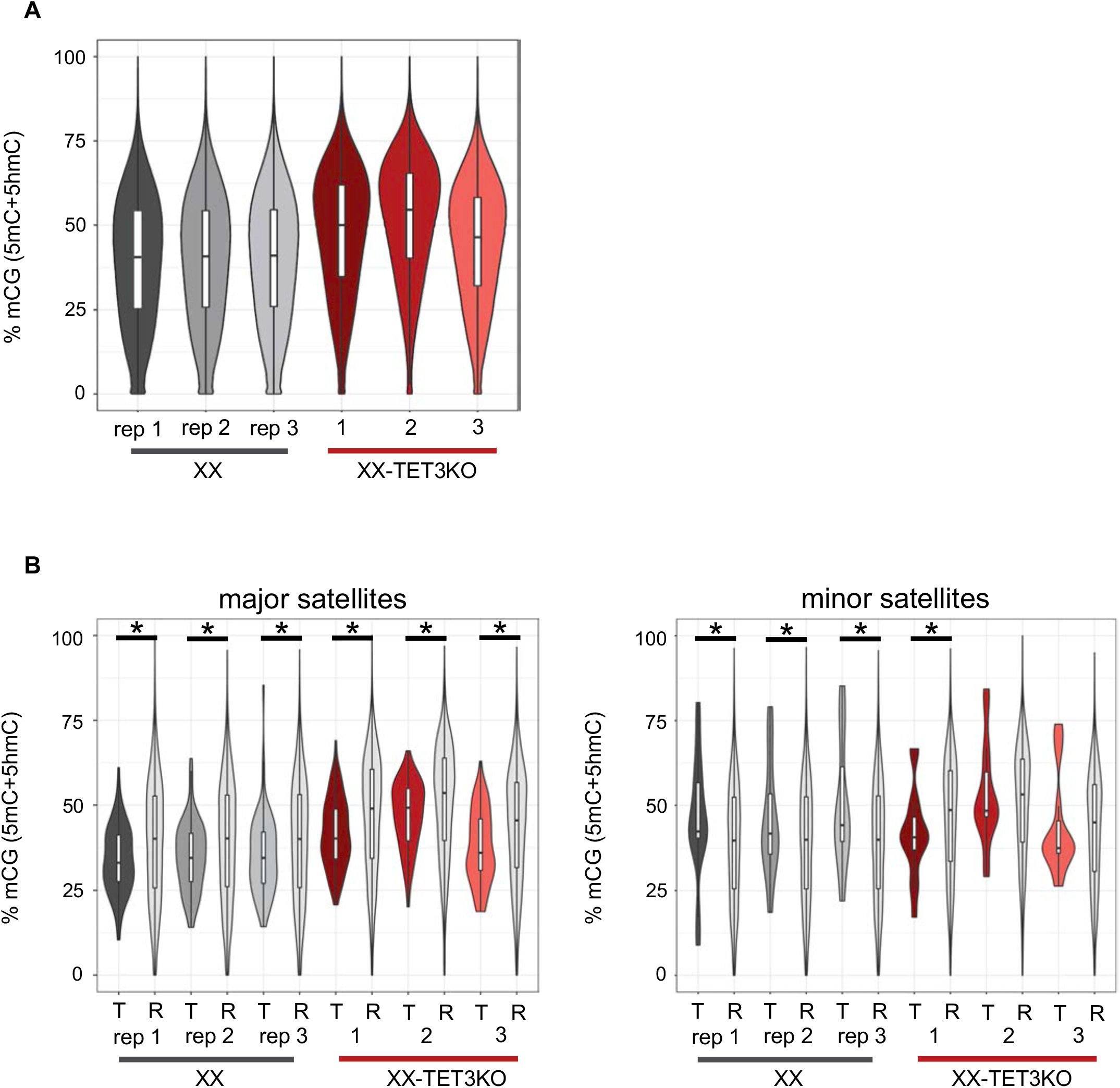
DNA methylation analysis of major and minor satellite repeats (A) Distribution of % CG methylation in each sample over 1kb windows tiled genome-wide. (B) Distribution of % CG methylation over major (left) and minor (right) T = satellites (UCSC repeatmasker) and R = 200x randomly shuffled sites, for each sample. * = significant difference by bootstrap testing, p <= 0.05 (one-tailed)

**Figure 5- figure supplement 1.**
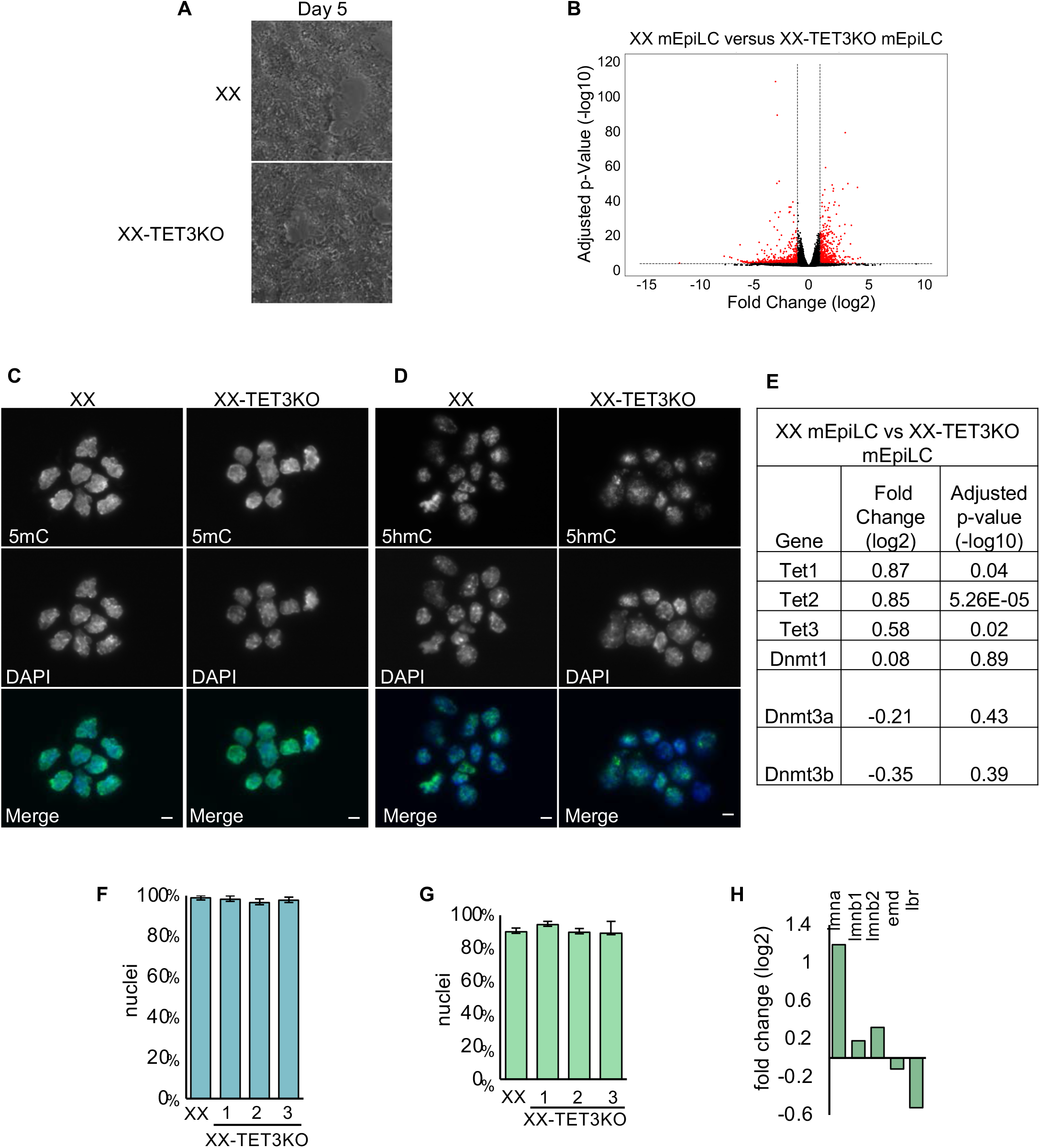
(A) Representative brightfield images of Day 5 mEpiLC differentiation in XX and XX-TET3KO cells. XX-TET3KO mEpiLICs exhibit considerably more cell death beginning day 4 of differentiation. A larger number of XX-TET3KO mEpiLCs were plated in order to visualize cell morphology in this image. (B) Full volcano plot comparing RNA-seq from XX and XX-TET3KO mEpiLCs showing genes excluded in Fig. 5A. Red dots indicate genes with fold changes (log2) < -1 or > 1 and adjusted p-value (-log10) <0.01. (C-D) IF of XX and XX-TET3KO 1 mEpiLCs using (C) 5mC and (D) 5hmC antibodies (green in Merge). DAPI (blue in Merge) indicates nuclei. White scale bar (Merge) indicates 5µm. (E) Table of fold change (log2) and adjusted p-values (-log10) of cytosine modification enzymes in XX/XX-TET3KO mEpiLCs. (F) Proportion of cells exhibiting H3K27me3 enrichment associated with Xist RNA in XX and XX-TET3KO mEpiLCs, n>100 cells/replicate. (G) Proportion of nuclei exhibiting H3K27me3 overlap with LMNB1 staining. Error bars indicate sd, n>100 cells/replicate. (H) Fold change (log2) of lamin-associated transcripts from XX mEpiLC/XX-TET3KO mEpiLC RNA-seq.

**Figure 7- figure supplement 1.**
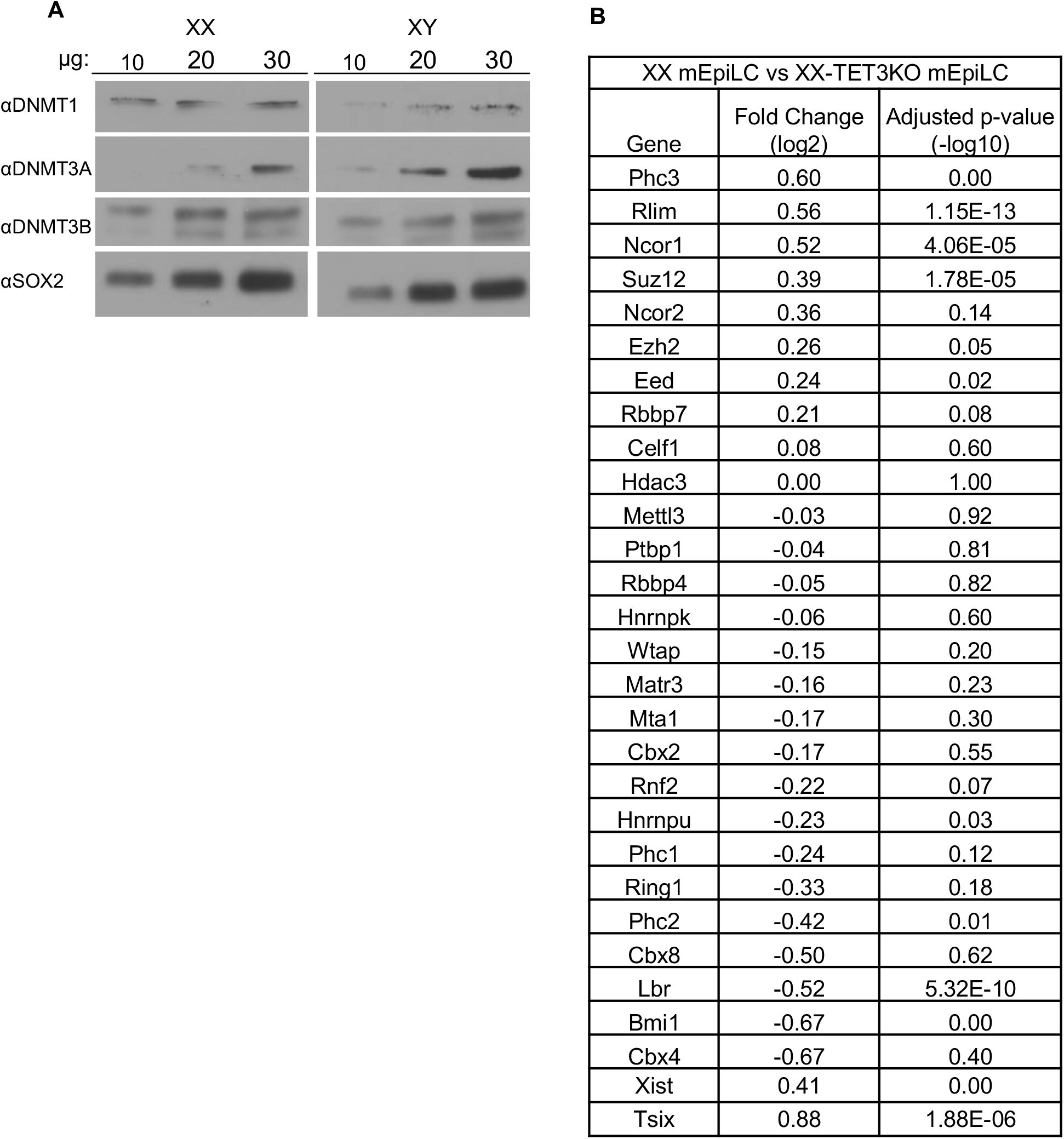
(A) DNMT1, DNMT3A, DNMT3B, and SOX2 immunoblots of increasing concentration of whole cell lysate prepared from XX and XY mESCs. (B) Table of fold change (log2) and adjusted p-values (-log10) of XIC-associated genes, comparing XX and XX-TET3KO mEpiLCs.

**Supplemental methods 1.**
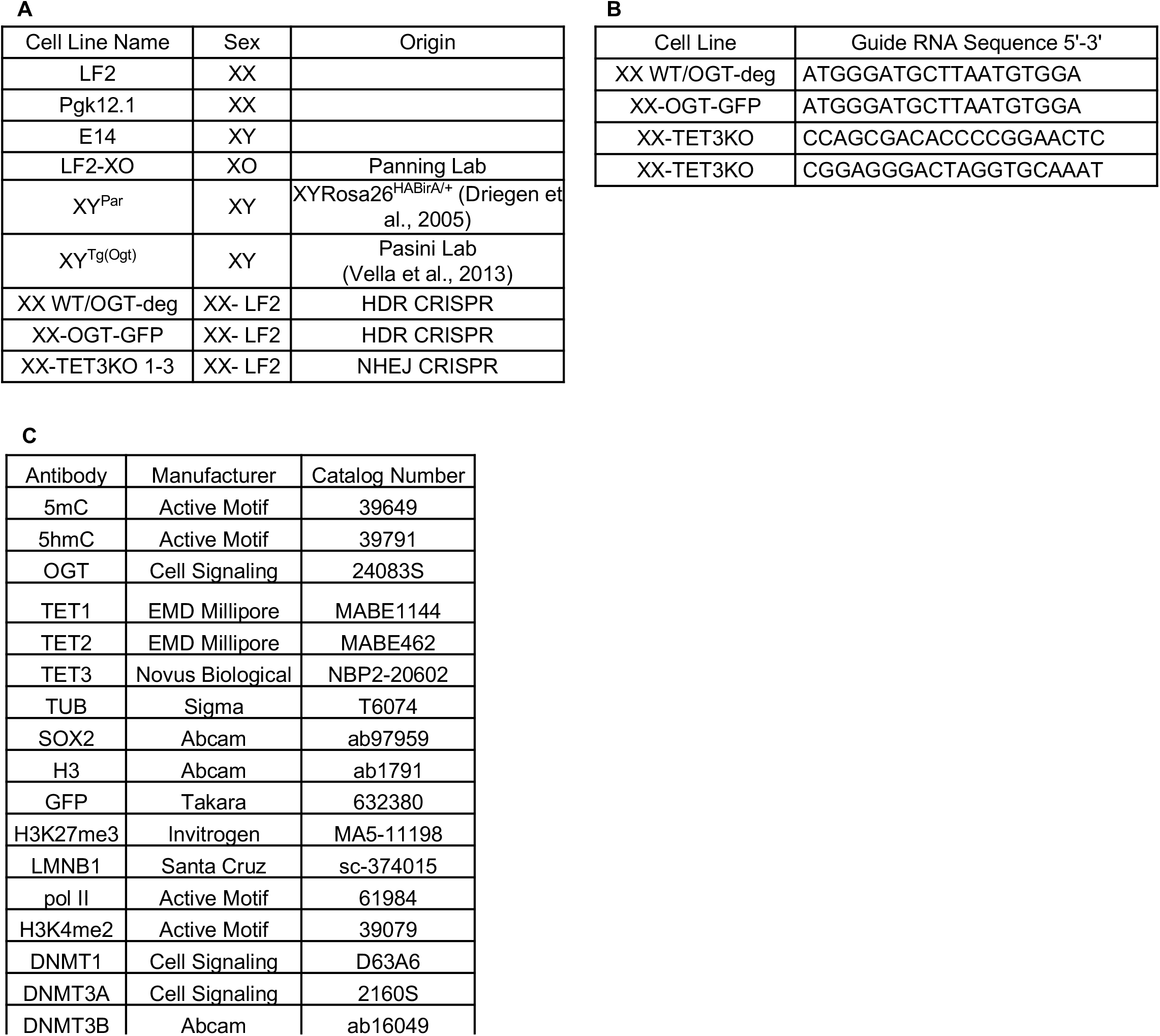
(A) Table of cell lines used and derived for experiments. (B) Table of guide RNA templates used for CRISPR/Cas9 gene editing. (C) Table of primary antibodies used for IF, immunoblots, and co-IPs.

**Supplemental methods 2.**
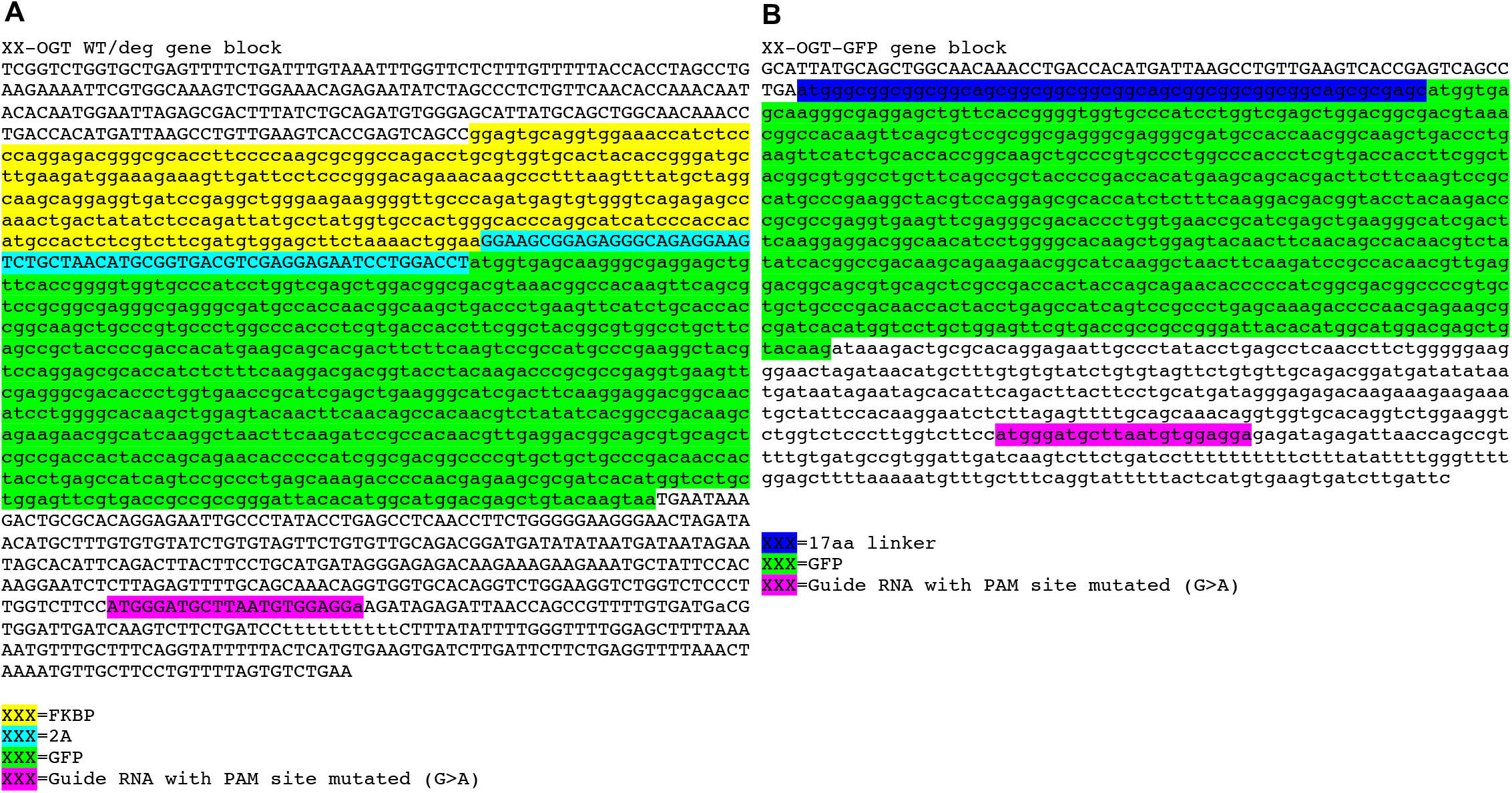
(A) Gene block of HDR insert for XX-OGT WT/deg cell line. (B) Gene block of HDR insert for XX-OGT-GFP cell line.

**Supplemental methods 3.**
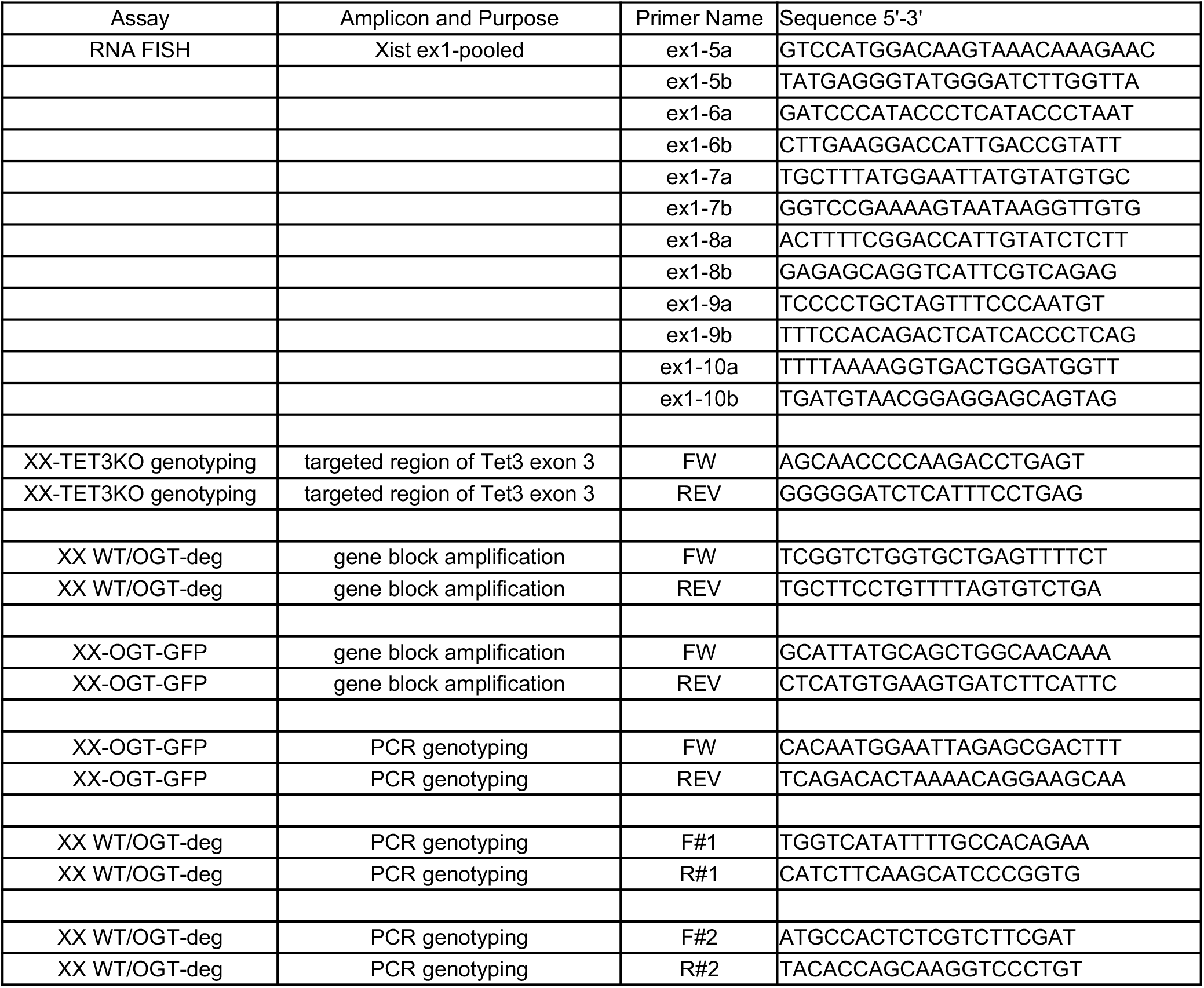
Table of primers used for creating the Xist RNA FISH probe, XX-TET3KO genotyping, XX WT/OGT-deg gene block amplification/PCR genotyping, and XX-OGT-GFP gene block amplification/PCR genotyping.

**Supplemental methods 4.**
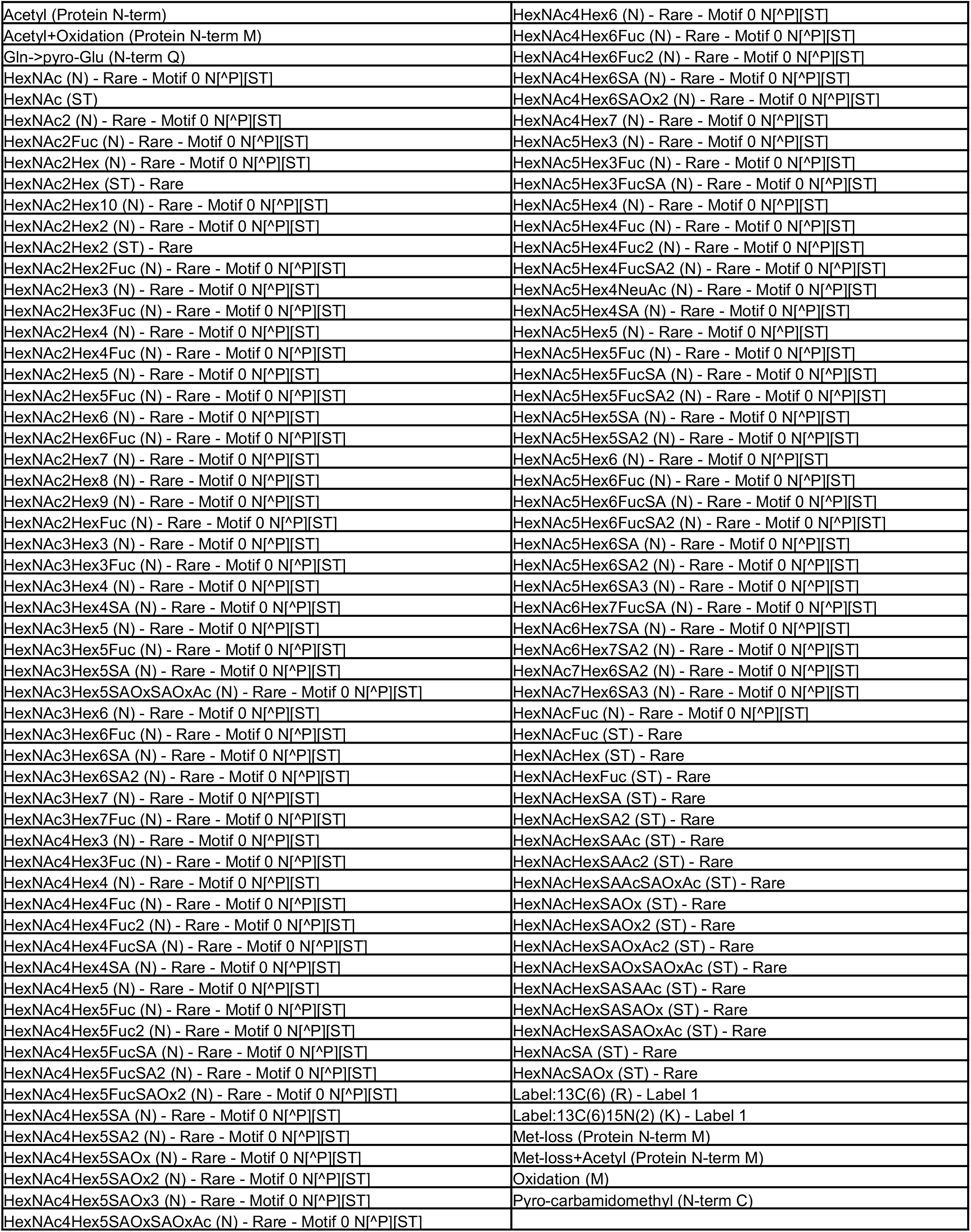
Table of variable modifications used in the mass spectrometry analysis.

**Supplemental methods 5.**
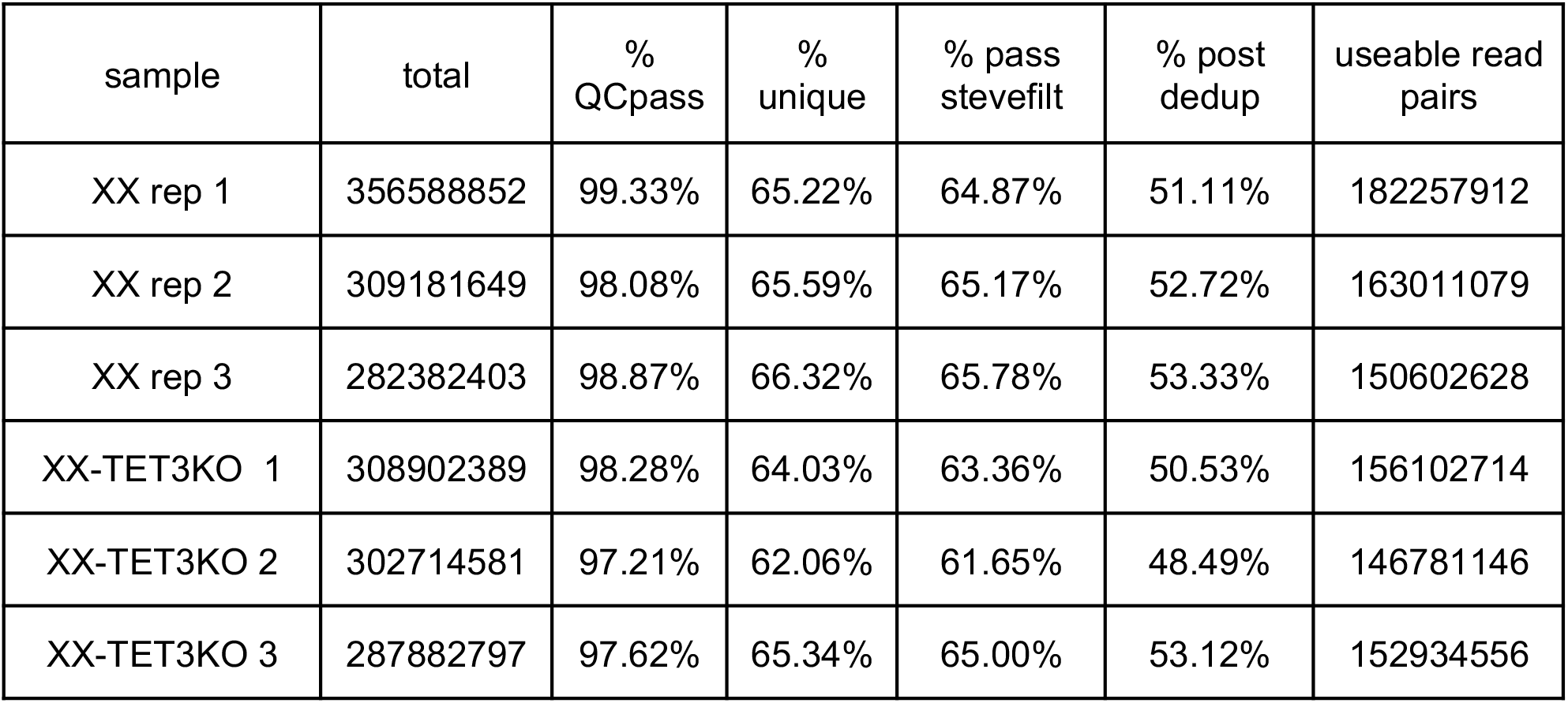
Table of Alignment statistics for WGBS libraries. Total = total number of read pairs obtained; % QCpass = percent of total remaining after QC filtering by trim_galore; % unique = percent of total that passed previous filters and aligned uniquely to mm10; % pass stevefilt = percent of total that passed previous filters and passed conversion filter (see methods); % post dedup = percent of total that passed previous filters and remained after PCR duplicates were removed; useable read pairs = final count of read pairs passing all filters that were used for downstream analyses.

## References

Baker, P. R., & Chalkley, R. J. (2014). MS-viewer: A web-based spectral viewer for proteomics results. Molecular & Cellular Proteomics: MCP, 13(5), 1392–1396. https://doi.org/10.1074/mcp.O113.037200

Baker, P. R., Trinidad, J. C., & Chalkley, R. J. (2011). Modification site localization scoring integrated into a search engine. Molecular & Cellular Proteomics: MCP, 10(7), M111.008078. https://doi.org/10.1074/mcp.M111.008078

Bauer, C., Göbel, K., Nagaraj, N., Colantuoni, C., Wang, M., Müller, U., Kremmer, E., Rottach, A., & Leonhardt, H. (2015). Phosphorylation of TET Proteins Is Regulated via O-GlcNAcylation by the O-Linked N-Acetylglucosamine Transferase (OGT). The Journal of Biological Chemistry, 290(8), 4801–4812. https://doi.org/10.1074/jbc.M114.605881

Benjamini Y, Hochberg Y. (1995) Controlling the False Discovery Rate: A Practical and Powerful Approach to Multiple Testing. Journal of the Royal Statistical Society. Series B (Methodological), 57(1), 289–300.

Boeren, J., & Gribnau, J. (2021). Xist-mediated chromatin changes that establish silencing of an entire X chromosome in mammals. Current Opinion in Cell Biology, 70, 44–50. https://doi.org/10.1016/j.ceb.2020.11.004

Chaumeil, J., Le Baccon, P., Wutz, A., & Heard, E. (2006). A novel role for Xist RNA in the formation of a repressive nuclear compartment into which genes are recruited when silenced. Genes & Development, 20(16), 2223–2237. https://doi.org/10.1101/gad.380906

Chen, Q., Chen, Y., Bian, C., Fujiki, R., & Yu, X. (2013). TET2 promotes histone O-GlcNAcylation during gene transcription. Nature, 493(7433), 561–564. https://doi.org/10.1038/nature11742

Choi, J., Clement, K., Huebner, A. J., Webster, J., Rose, C. M., Brumbaugh, J., Walsh, R. M., Lee, S., Savol, A., Etchegaray, J.-P., Gu, H., Boyle, P., Elling, U., Mostoslavsky, R., Sadreyev, R., Park, P. J., Gygi, S. P., Meissner, A., & Hochedlinger, K. (2017). DUSP9 Modulates DNA Hypomethylation in Female Mouse Pluripotent Stem Cells. Cell Stem Cell, 20(5), 706–719.e7. https://doi.org/10.1016/j.stem.2017.03.002

Cline, T. W., & Meyer, B. J. (1996). Vive la différence: Males vs females in flies vs worms. Annual Review of Genetics, 30, 637–702. https://doi.org/10.1146/annurev.genet.30.1.637

Dai, H.-Q., Wang, B.-A., Yang, L., Chen, J.-J., Zhu, G.-C., Sun, M.-L., Ge, H., Wang, R., Chapman, D. L., Tang, F., Sun, X., & Xu, G.-L. (2016). TET-mediated DNA demethylation controls gastrulation by regulating Lefty-Nodal signalling. Nature, 538(7626), 528–532. https://doi.org/10.1038/nature20095

Dawlaty, M. M., Breiling, A., Le, T., Barrasa, M. I., Raddatz, G., Gao, Q., Powell, B. E., Cheng, A. W., Faull, K. F., Lyko, F., & Jaenisch, R. (2014). Loss of Tet enzymes compromises proper differentiation of embryonic stem cells. Developmental Cell, 29(1), 102–111. https://doi.org/10.1016/j.devcel.2014.03.003

Driegen, S., Ferreira, R., van Zon, A., Strouboulis, J., Jaegle, M., Grosveld, F., Philipsen, S., & Meijer, D. (2005). A generic tool for biotinylation of tagged proteins in transgenic mice. Transgenic Research, 14(4), 477–482. https://doi.org/10.1007/s11248-005-7220-2

Gu, T.-P., Guo, F., Yang, H., Wu, H.-P., Xu, G.-F., Liu, W., Xie, Z.-G., Shi, L., He, X., Jin, S., Iqbal, K., Shi, Y. G., Deng, Z., Szabó, P. E., Pfeifer, G. P., Li, J., & Xu, G.-L. (2011). The role of Tet3 DNA dioxygenase in epigenetic reprogramming by oocytes. Nature, 477(7366), 606–610. https://doi.org/10.1038/nature10443

Habibi, E., Brinkman, A. B., Arand, J., Kroeze, L. I., Kerstens, H. H. D., Matarese, F., Lepikhov, K., Gut, M., Brun-Heath, I., Hubner, N. C., Benedetti, R., Altucci, L., Jansen, J. H., Walter, J., Gut, I. G., Marks, H., & Stunnenberg, H. G. (2013). Whole-Genome Bisulfite Sequencing of Two Distinct Interconvertible DNA Methylomes of Mouse Embryonic Stem Cells. Cell Stem Cell, 13(3), 360–369. https://doi.org/10.1016/j.stem.2013.06.002

Halim, A., Westerlind, U., Pett, C., Schorlemer, M., Rüetschi, U., Brinkmalm, G., Sihlbom, C., Lengqvist, J., Larson, G., & Nilsson, J. (2014). Assignment of Saccharide Identities through Analysis of Oxonium Ion Fragmentation Profiles in LC–MS/MS of Glycopeptides. Journal of Proteome Research, 13(12), 6024–6032. https://doi.org/10.1021/pr500898r

Hall, L. L., & Lawrence, J. B. (2010). XIST RNA and Architecture of the Inactive X chromosome: Implications for the Repeat Genome. Cold Spring Harbor Symposia on Quantitative Biology, 75, 345–356. https://doi.org/10.1101/sqb.2010.75.030

Hrit, J., Goodrich, L., Li, C., Wang, B.-A., Nie, J., Cui, X., Martin, E. A., Simental, E., Fernandez, J., Liu, M. Y., Nery, J. R., Castanon, R., Kohli, R. M., Tretyakova, N., He, C., Ecker, J. R., Goll, M., & Panning, B. (2018). OGT binds a conserved C-terminal domain of TET1 to regulate TET1 activity and function in development. ELife, 7, e34870. https://doi.org/10.7554/eLife.34870

Ito, R., Katsura, S., Shimada, H., Tsuchiya, H., Hada, M., Okumura, T., Sugawara, A., & Yokoyama, A. (2014). TET3-OGT interaction increases the stability and the presence of OGT in chromatin. Genes to Cells: Devoted to Molecular & Cellular Mechanisms, 19(1), 52–65. https://doi.org/10.1111/gtc.12107

Khoueiry, R., Sohni, A., Thienpont, B., Luo, X., Velde, J. V., Bartoccetti, M., Boeckx, B., Zwijsen, A., Rao, A., Lambrechts, D., & Koh, K. P. (2017). Lineage-specific functions of TET1 in the postimplantation mouse embryo. Nature Genetics, 49(7), 1061–1072. https://doi.org/10.1038/ng.3868

Kohlmaier, A., Savarese, F., Lachner, M., Martens, J., Jenuwein, T., & Wutz, A. (2004). A Chromosomal Memory Triggered by Xist Regulates Histone Methylation in X Inactivation. PLOS Biology, 2(7), e171. https://doi.org/10.1371/journal.pbio.0020171

Kreppel, L. K., Blomberg, M. A., & Hart, G. W. (1997). Dynamic Glycosylation of Nuclear and Cytosolic Proteins: CLONING AND CHARACTERIZATION OF A UNIQUE O-GlcNAc TRANSFERASE WITH MULTIPLE TETRATRICOPEPTIDE REPEATS *. Journal of Biological Chemistry, 272(14), 9308–9315. https://doi.org/10.1074/jbc.272.14.9308

Krueger, F. (2021). FelixKrueger/TrimGalore [Perl]. https://github.com/FelixKrueger/TrimGalore (Original work published 2016)

Krueger, F., & Andrews, S. R. (2011). BKrueger, F. (2021). FelixKrueger/TrimGalore [Perl]. https://github.com/FelixKrueger/TrimGalore (Original work published 2016)

Lewis, B. A., & Hanover, J. A. (2014). O-GlcNAc and the Epigenetic Regulation of Gene Expression. The Journal of Biological Chemistry, 289(50), 34440–34448. https://doi.org/10.1074/jbc.R114.595439

Maynard, J., & Chalkley, R. J. (2020). Methods for Enrichment and Assignment of N-Acetylglucosamine Modification Sites. Molecular & Cellular Proteomics: MCP, 20, 100031. https://doi.org/10.1074/mcp.R120.002206

Mlynarczyk-Evans, S., Royce-Tolland, M., Alexander, M. K., Andersen, A. A., Kalantry, S., Gribnau, J., & Panning, B. (2006). X Chromosomes Alternate between Two States prior to Random X-Inactivation. PLoS Biology, 4(6). https://doi.org/10.1371/journal.pbio.0040159

Myers, S. A., Panning, B., & Burlingame, A. L. (2011). Polycomb repressive complex 2 is necessary for the normal site-specific O-GlcNAc distribution in mouse embryonic stem cells. Proceedings of the National Academy of Sciences, 108(23), 9490–9495. https://doi.org/10.1073/pnas.1019289108.

Nabet, B., Roberts, J. M., Buckley, D. L., Paulk, J., Dastjerdi, S., Yang, A., Leggett, A. L., Erb, M. A., Lawlor, M. A., Souza, A., Scott, T. G., Vittori, S., Perry, J. A., Qi, J., Winter, G. E., Wong, K.-K., Gray, N. S., & Bradner, J. E. (2018). The dTAG system for immediate and target-specific protein degradation. Nature Chemical Biology, 14(5), 431–441. https://doi.org/10.1038/s41589-018-0021-8

Pandya-Jones, A., Markaki, Y., Serizay, J., Chitiashvili, T., Mancia, W., Damianov, A., Chronis, C., Papp, B., Chen, C.-K., McKee, R., Wang, X.-J., Chau, A., Sabri, S., Leonhardt, H., Zheng, S., Guttman, M., Black, Douglas. L., & Plath, K. (2020). A protein assembly mediates Xist localization and gene silencing. Nature, 587(7832), 145–151. https://doi.org/10.1038/s41586-020-2703-0

Panning, B., & Jaenisch, R. (1996). DNA hypomethylation can activate Xist expression and silence X-linked genes. Genes & Development, 10(16), 1991–2002. https://doi.org/10.1101/gad.10.16.1991

Pantier, R., Tatar, T., Colby, D., & Chambers, I. (2019). Endogenous epitope-tagging of Tet1, Tet2 and Tet3 identifies TET2 as a naïve pluripotency marker. Life Science Alliance, 2(5). https://doi.org/10.26508/lsa.201900516

Parry, A., Rulands, S., & Reik, W. (2021). Active turnover of DNA methylation during cell fate decisions. Nature Reviews Genetics, 22(1), 59–66. https://doi.org/10.1038/s41576-020-00287-8

Pignatta, D., Bell, G. W., & Gehring, M. (2015). Whole Genome Bisulfite Sequencing and DNA Methylation Analysis from Plant Tissue. Bio-Protocol, 5(4), e1407–e1407.

Plath, K., Mlynarczyk-Evans, S., Nusinow, D. A., & Panning, B. (2002). Xist RNA and the Mechanism of X Chromosome Inactivation. Annual Review of Genetics, 36(1), 233–278. https://doi.org/10.1146/annurev.genet.36.042902.092433

Ran, F. A., Hsu, P. D., Wright, J., Agarwala, V., Scott, D. A., & Zhang, F. (2013). Genome engineering using the CRISPR-Cas9 system. Nature Protocols, 8(11), 2281–2308. https://doi.org/10.1038/nprot.2013.143

Rego, A., Sinclair, P. B., Tao, W., Kireev, I., & Belmont, A. S. (2008). The facultative heterochromatin of the inactive X chromosome has a distinctive condensed ultrastructure. Journal of Cell Science, 121(Pt 7), 1119–1127. https://doi.org/10.1242/jcs.026104

Reik, W., & Surani, M. A. (2015). Germline and Pluripotent Stem Cells. Cold Spring Harbor Perspectives in Biology, 7(11). https://doi.org/10.1101/cshperspect.a019422

Ross, S. E., & Bogdanovic, O. (2019). TET enzymes, DNA demethylation and pluripotency. Biochemical Society Transactions, 47(3), 875–885. https://doi.org/10.1042/BST20180606

R Core Team. (2017). R: A language and environment for statistical computing. R Foundation for Statistical Computing, Vienna, Austria. https://www.R-project.org/

Sarma, K., Cifuentes-Rojas, C., Ergun, A., del Rosario, A., Jeon, Y., White, F., Sadreyev, R., & Lee, J. T. (2014). ATRX Directs Binding of PRC2 to Xist RNA and Polycomb Targets. Cell, 159(4), 869–883. https://doi.org/10.1016/j.cell.2014.10.019

Schulz, E. G., Meisig, J., Nakamura, T., Okamoto, I., Sieber, A., Picard, C., Borensztein, M., Saitou, M., Blüthgen, N., & Heard, E. (2014). The Two Active X Chromosomes in Female ESCs Block Exit from the Pluripotent State by Modulating the ESC Signaling Network. Cell Stem Cell, 14(2), 203–216. https://doi.org/10.1016/j.stem.2013.11.022

Seo, H. G., Kim, H. B., Kang, M. J., Ryum, J. H., Yi, E. C., & Cho, J. W. (2016). Identification of the nuclear localisation signal of O -GlcNAc transferase and its nuclear import regulation. Scientific Reports, 6(1), 34614. https://doi.org/10.1038/srep34614

Shafi, R., Iyer, S. P. N., Ellies, L. G., O’Donnell, N., Marek, K. W., Chui, D., Hart, G. W., & Marth, J. D. (2000). The O-GlcNAc transferase gene resides on the X chromosome and is essential for embryonic stem cell viability and mouse ontogeny. Proceedings of the National Academy of Sciences, 97(11), 5735–5739. https://doi.org/10.1073/pnas.100471497

Shi, F.-T., Kim, H., Lu, W., He, Q., Liu, D., Goodell, M. A., Wan, M., & Songyang, Z. (2013). Ten-Eleven Translocation 1 (Tet1) Is Regulated by O-Linked N-Acetylglucosamine Transferase (Ogt) for Target Gene Repression in Mouse Embryonic Stem Cells. The Journal of Biological Chemistry, 288(29), 20776–20784. https://doi.org/10.1074/jbc.M113.460386

Smit A, Hubley R, Green P. (1996) RepeatMasker Open-3.0, http://www.repeatmasker.org/

Song, J., Janiszewski, A., De Geest, N., Vanheer, L., Talon, I., El Bakkali, M., Oh, T., & Pasque, V. (2019). X-Chromosome Dosage Modulates Multiple Molecular and Cellular Properties of Mouse Pluripotent Stem Cells Independently of Global DNA Methylation Levels. Stem Cell Reports, 12(2), 333–350. https://doi.org/10.1016/j.stemcr.2018.12.004

Takahashi, S., Kobayashi, S., & Hiratani, I. (2018). Epigenetic differences between naïve and primed pluripotent stem cells. Cellular and Molecular Life Sciences, 75(7), 1191–1203. https://doi.org/10.1007/s00018-017-2703-x

Trinidad, J. C., Barkan, D. T., Gulledge, B. F., Thalhammer, A., Sali, A., Schoepfer, R., & Burlingame, A. L. (2012). Global Identification and Characterization of Both O-GlcNAcylation and Phosphorylation at the Murine Synapse. Molecular & Cellular Proteomics : MCP, 11(8), 215–229. https://doi.org/10.1074/mcp.O112.018366

von Meyenn, F., Iurlaro, M., Habibi, E., Liu, N. Q., Salehzadeh-Yazdi, A., Santos, F., Petrini, E., Milagre, I., Yu, M., Xie, Z., Kroeze, L. I., Nesterova, T. B., Jansen, J. H., Xie, H., He, C., Reik, W., & Stunnenberg, H. G. (2016). Impairment of DNA Methylation Maintenance Is the Main Cause of Global Demethylation in Naive Embryonic Stem Cells. Molecular Cell, 62(6), 848–861. https://doi.org/10.1016/j.molcel.2016.04.025

Wickham H. (2016) Ggplot2: Elegant Graphics for Data Analysis. Springer-Verlag New York;https://ggplot2.tidyverse.org

Wijchers, P. J., Geeven, G., Eyres, M., Bergsma, A. J., Janssen, M., Verstegen, M., Zhu, Y., Schell, Y., Vermeulen, C., Wit, E. de, & Laat, W. de. (2015). Characterization and dynamics of pericentromere-associated domains in mice. Genome Research. https://doi.org/10.1101/gr.186643.114

Wutz, A., & Jaenisch, R. (2000). A Shift from Reversible to Irreversible X Inactivation Is Triggered during ES Cell Differentiation. Molecular Cell, 5(4), 695–705. https://doi.org/10.1016/S1097-2765(00)80248-8

Wutz, A., Rasmussen, T. P., & Jaenisch, R. (2002). Chromosomal silencing and localization are mediated by different domains of Xist RNA. Nature Genetics, 30(2), 167–174. https://doi.org/10.1038/ng820

Xiao, P., Zhou, X., Zhang, H., Xiong, K., Teng, Y., Huang, X., Cao, R., Wang, Y., & Liu, H. (2013). Characterization of the nuclear localization signal of the mouse TET3 protein. Biochemical and Biophysical Research Communications, 439(3), 373–377. https://doi.org/10.1016/j.bbrc.2013.08.075

Zhang, Q., Liu, X., Gao, W., Li, P., Hou, J., Li, J., & Wong, J. (2014). Differential Regulation of the Ten-Eleven Translocation (TET) Family of Dioxygenases by O-Linked β-N-Acetylglucosamine Transferase (OGT). The Journal of Biological Chemistry, 289(9), 5986–5996. https://doi.org/10.1074/jbc.M113.524140

Zvetkova, I., Apedaile, A., Ramsahoye, B., Mermoud, J. E., Crompton, L. A., John, R., Feil, R., & Brockdorff, N. (2005). Global hypomethylation of the genome in XX embryonic stem cells. Nature Genetics, 37(11), 1274–1279. https://doi.org/10.1038/ng1663

